# Hippocampal neuronal activity is aligned with action plans

**DOI:** 10.1101/2024.09.05.611533

**Authors:** Ipshita Zutshi, Athina Apostolelli, Wannan Yang, Zheyang (Sam) Zheng, Tora Dohi, Edoardo Balzani, Alex H Williams, Cristina Savin, György Buzsáki

**Affiliations:** Neuroscience Institute, New York University Grossman School of Medicine; Center for Neural Science, New York University; Center for Data Science, New York University; Center for Computational Neuroscience, Flatiron Institute

## Abstract

Neurons in the hippocampus are correlated with different variables, including space, time, sensory cues, rewards, and actions, where the extent of tuning depends on ongoing task demands. However, it remains uncertain whether such diverse tuning corresponds to distinct functions within the hippocampal network or if a more generic computation can account for these observations. To disentangle the contribution of externally driven cues versus internal computation, we developed a task in mice where space, auditory tones, rewards, and context were juxtaposed with changing relevance. High-density electrophysiological recordings revealed that neurons were tuned to each of these modalities. By comparing movement paths and action sequences, we observed that external variables had limited direct influence on hippocampal firing. Instead, spiking was influenced by online action plans modulated by goal uncertainty. Our results suggest that internally generated cell assembly sequences are selected and updated by action plans toward deliberate goals. The apparent tuning of hippocampal neuronal spiking to different sensory modalities might emerge due to alignment to the afforded action progression within a task rather than representation of external cues.

## INTRODUCTION

The hippocampus is often portrayed as a structure at the top of the cortical hierarchy, which receives highly processed sensory modalities(Felleman and Van Essen, 1991). It is assumed to synthesize those modalities into abstract space and time, explicitly reflected by its neurons’ place and time fields(Breathnach, 1980; Eichenbaum, 2000; Manns et al., 2007). Other considerations suggest that it is part of a top-down effector system involved in planning and initiating voluntary actions(Arezzo and Vaughan, 1975; Foster et al., 1989; Glass, 2019; Halgren, 1991; von Holst, 1954; Miller et al., 2017; Vanderwolf, 1969; Wiener and Korshunov, 1995). In collaboration with frontal cortical areas(Fried et al., 2017; Miller and Cohen, 2001; Numan, 1978, 2015), it can also sustain working memory, possibly related to the preparation for upcoming actions(Fuster, 1991; Gaesser et al., 2013; Olton et al., 1980). Combining the sensory synthesis and effector systems supports effective navigation in an environment(Mcnaughton et al., 1996) in the form of a predictive code(Stachenfeld et al., 2017). By disengaging from the environment, the hippocampus can support memory and planning (Buzsáki and Moser, 2013).

These views have been supported by carefully performed experiments, of which spatial navigation, particularly the sensory synthesis view, has received the most prominent attention (Eichenbaum, 2017; Lisman et al., 2017; Schiller et al., 2015). Yet, whether one framework can satisfactorily incorporate competing views and explain experimental data from all domains has remained a challenge(O’Keefe and Krupic, 2021). Part of the problem is that the experimental design strongly influences the findings and their interpretation(Han et al., 2023). By designing experiments where space, time, action, memory, or sensory modality is the relevant domain, one finds a description of place fields, time fields, reward cells, speed cells, tone cells, choice cells, or “engram” cells in the hippocampus(Allen et al., 2016; Aronov et al., 2017; Breathnach, 1980; Fischler-Ruiz et al., 2021; Gauthier and Tank, 2018; Josselyn and Tonegawa, 2020; MacDonald et al., 2011; Manns et al., 2007; Purandare and Mehta, 2023; Radvansky et al., 2021; Sun et al., 2020; Tuncdemir et al., 2022). The implicit assumption from these results is that each label reflects unique computations performed by the hippocampus, generating “representations” specific to the relevant modality and reflecting different functions of hippocampal firing. Yet, a common confound is that the behavior of animals becomes inevitably linked to the relevant domain(Hasnain et al., 2023; Vanderwolf, 1969).

To disentangle the various sensory and task-specific variables, motor actions and intentions, we designed a task that incorporated multiple behaviorally relevant domains with careful controls. The design allowed quantifying the relationship between hippocampal neuronal firing patterns and multiple juxtaposed variables (including space, relevant and non-relevant auditory tones, numerous choices, rewards, context, and actions) in the same experiment. By comparing similar movement trajectories and actions across task contingencies, we tested whether the firing of hippocampal neurons was driven by a general internal computation applied across all modalities.

## RESULTS

### Navigation task with juxtaposed spatial coordinates and self-controlled auditory tones

We implemented an agent-controlled auditory navigation task that could disambiguate tuning to spatial coordinates versus auditory tones (**Fig. 1a, Extended Data Fig. 1a, Movies 1,2**). Mice (*n = 6*) ran on a linear track with 7 equally spaced water ports. During NO-TONE (NT) trials, they ran back and forth to collect water rewards at either end with no auditory tones playing. During TONE trials, a tone ascended in frequency (2-25kHz) in a closed-loop manner controlled by the mouse’s spatial location. The sweep gain between frequency and position was randomized from trial to trial so that the same frequency occurred at different spatial locations across trials (see Methods). Mice must attend to the tone and lick at the closest water port whenever the target frequency (22 kHz) is reached. Return runs had no tone associated with them and served as control. Incorrect licks would lead to cessation of the tone and require the mouse to run back to the home port to initiate a new trial. Thus, during TONE trials, spatial (room) coordinates were uninformative because tones were the only predictor of reward location. Mice learned the task within three weeks and reached ∼70% performance (**Fig. 1b-d, Extended Data Fig. 1b-c**). They performed better at the nearest target port, but the performance was roughly equal across the other ports (**Fig. 1e**). Most errors involved the mice licking one port sooner than the target, and very few overshoots occurred beyond the target port (**Extended Data Fig. 1d-e**).

To confirm that mice attended to the auditory tones throughout the run, we implemented probe trials in two mice. Here, the tones would abruptly truncate at either 4, 8, or 16kHz, allowing us to estimate when the mice could successfully extrapolate the slope of the frequency change and predict the target port (**Fig. 1g**). Probe trials were randomly interspersed with tone trials at a likelihood of ∼27%. While performance deteriorated at 4 and 8kHz, 16 kHz probe trials were similar to controls, though there was variation in this performance across different ports (**Fig. 1h, Extended Data Fig. 1f-h**). These results suggest that, on average, mice could estimate and plan their approach to the target port sometime before 16 kHz, but after 8 kHz tones.

**Figure 1:**
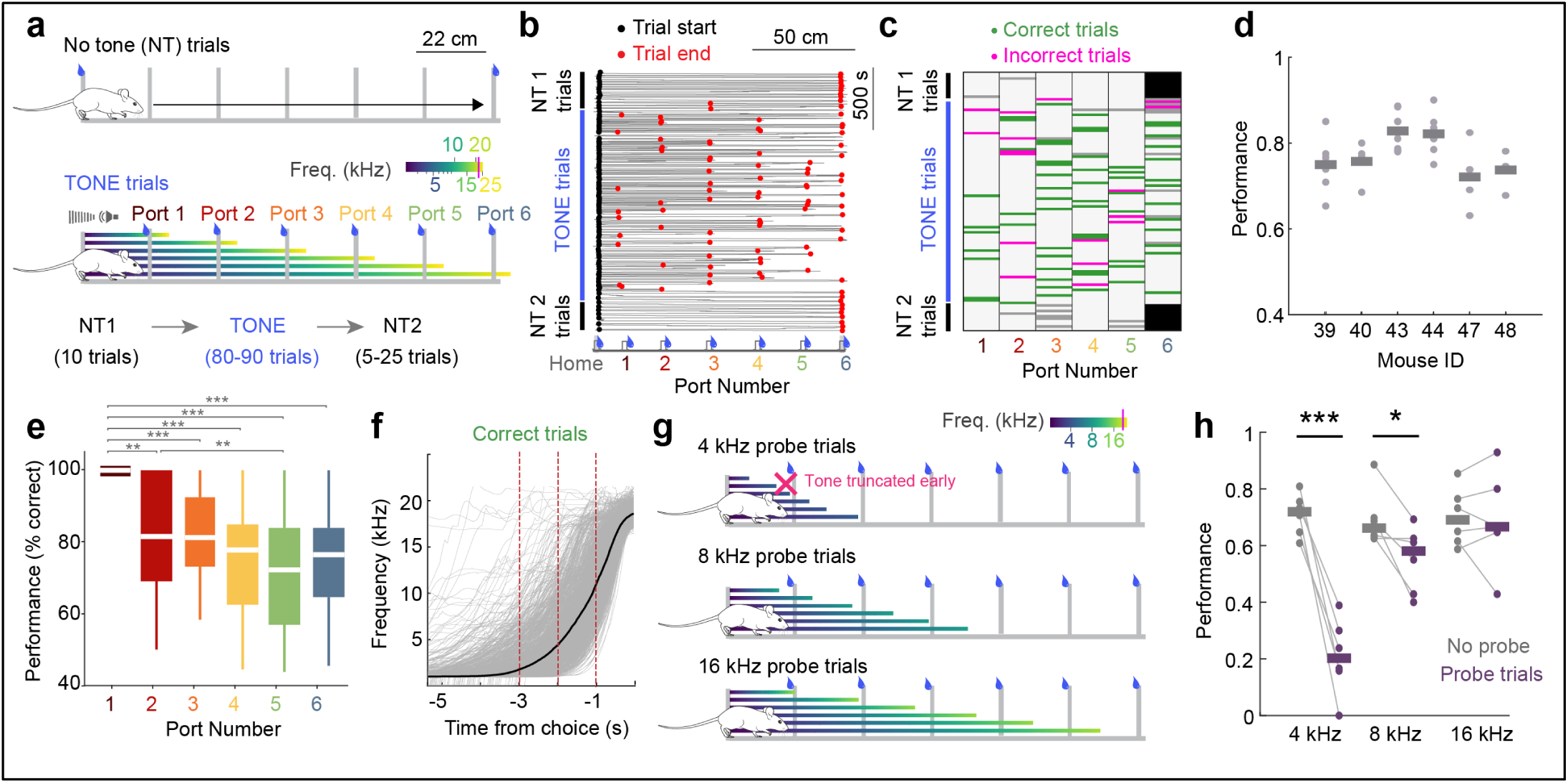
Agency-controlled auditory navigation task to dissociate non-spatial and spatial features. (a) Schematic of the task. TONE trials were preceded and followed by blocks of no-tone (NT) trials. During NT trials mice ran back and forth to receive a water reward at either end with no tones. During TONE trials, tone frequency was controlled by the mouse’s spatial location in a closed-loop manner and predicted the reward location. Gradient lines (*blue to yellow;* frequency sweep; 0-25 kHz) during TONE trials correspond to individual trial types targeting ports 1-6 (target frequency = 22 kHz, red line in the colormap). Return runs to home port had no tones playing. (b) Trajectory of a mouse from an example session. Mice ran to either end of the track during NT trials. During TONE trials, mice would either run straight back to the home port after making a choice or sample successive ports before returning. *Black circles*, trial start; *red circles*, trial end (see Movies 1 & 2). (c) Heatmap shows licks detected at each port (*black,* NT licks; *green,* TONE correct licks*; magenta,* TONE incorrect licks*; gray,* other licks, generally occurring at middle ports during NT trials or after incorrect licks on TONE trials). (d) Average performance, i.e., fraction of correct trials, of included sessions (*line*, median, *dots*, individual sessions) from six mice. (e) Average performance, i.e., percentage of correct trials compared to all trials where a specific port was assigned the target. Mice performed best for the first port (n = 37 sessions from 6 mice, Friedman test followed by Tukey-Kramer post-hoc tests, Chi-sq (5,175) = 64.31, *p = 1.55 x 10*^-12^). (f) Mouse-controlled tone frequency progression of each trial (*gray lines*) in time. Average frequency for correct trials is superimposed (*black*). *Red lines* mark -3, -2, and -1s before the first lick. (g) Schematic showing random probe trials that were interspersed with TONE trials. Frequencies were cut off at 4, 8, or 16 kHz during probe trials. Within a session, only one frequency cutoff was used. (h) Performance during probe trials compared to non-probe trials within a session. Performance was impaired during 4 and 8 kHz probe trials and not affected during 16 kHz trials (Paired t-test, n = 6 sessions from 2 mice, 4 kHz: t(5) = 8.3014, *p = 4.14 x 10^-4^;* 8 kHz: t(5) = 3.0758, *p = 0.0276;* 16 kHz: t(5) = 0.2694, *p = 0.7984*). *\*p<0.05, **p<0.01, ***p<0.001.* All box plots show median ± interquartile; whiskers show range excluding outliers.

### Conjunctive non-spatial and spatial firing led to mixed hippocampal cell assemblies

Mice were implanted with 128-site double-sided silicon probes to maximize unit yield from the CA1 pyramidal cell layer (Huszár et al., 2022). We recorded a median of 235 neurons per session (83-372 cells; 8113 cells total with n = 6220 pyramidal cells and n = 1893 putative interneurons out of 37 sessions in 6 mice). We first examined the activity dynamics of individual neurons as mice were solving the task. There were clear examples of cells whose discharge ramped up (**Fig. 2a**, Cell 1) or down (**Fig. 2a**, Cell 2) prior to the approach to any water port during TONE trials. For descriptive purposes only, these neurons are referred to as “tone cells” based on a detected field in frequency space (see Methods(Aronov et al., 2017)). In addition to this non-spatial frequency field, several cells displayed combinations of tone responses and a separate spatial place field (**Fig. 2a**, Cell 3). While “tone cell” fields tiled the entire frequency landscape, there was a general overrepresentation of frequencies near the 22 kHz target (**Fig. 2b**). Tone cells constituted a significant fraction of all recorded CA1 cells (∼24% of a median of 132 active pyramidal neurons per session; a total of 1129 cells; **Extended Data Fig. 2a, b**). While many tone cells (such as Cell 1) tended to have low firing rates during NT trials, ∼50% of these cells also had stable place fields during NT trials (**Extended Data Fig. 2c, d**). During error trials, we observed that the firing of tone cells was driven by the mouse’s approach to its chosen port rather than the assigned target for the trial (**Extended Data Fig. 2e-g**), suggesting that such firing might reflect activity ramping up to the selected port (i.e., the mouse’s *progression to the chosen port*) rather than auditory frequencies.

**Figure 2:**
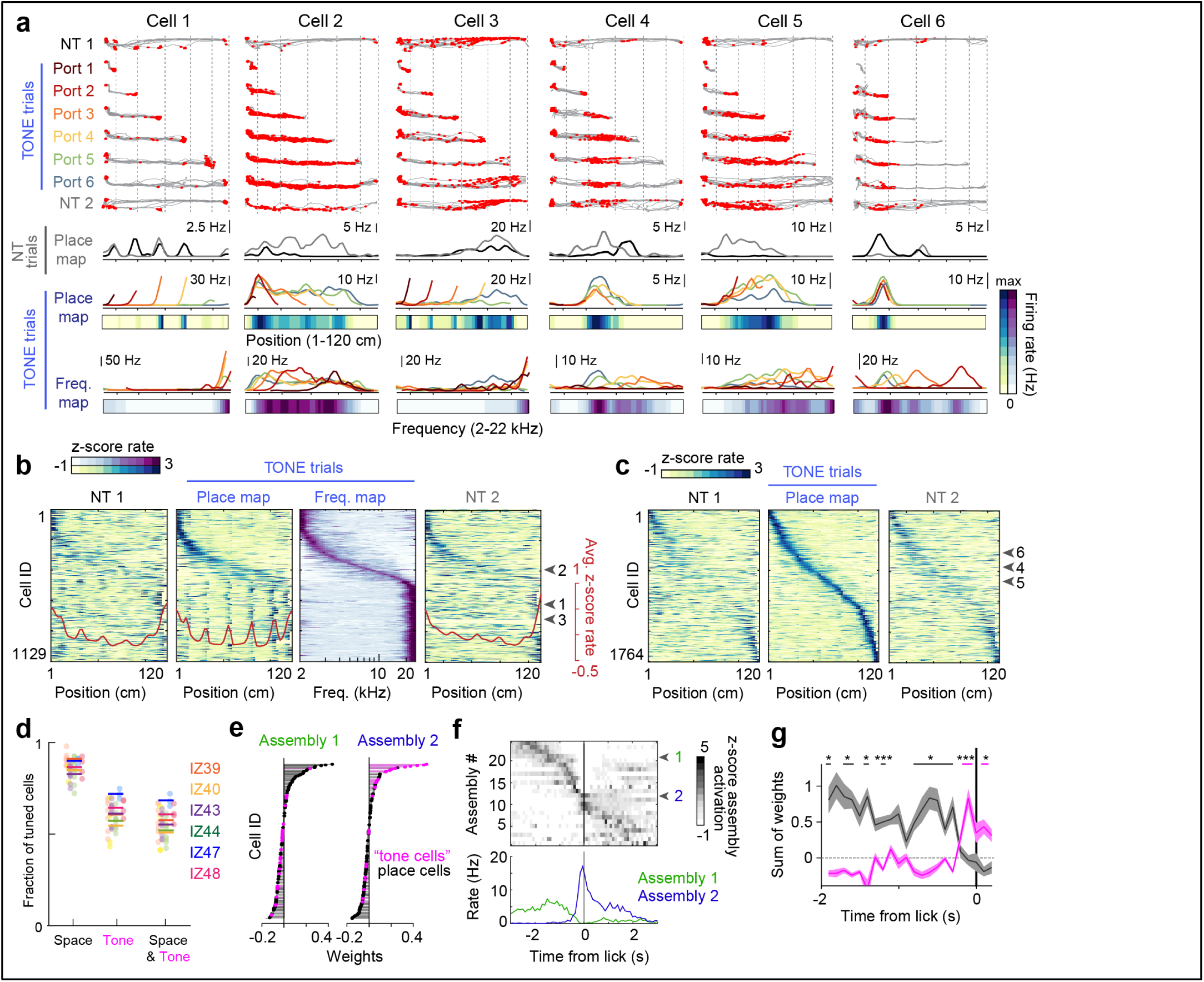
Conjunctive non-spatial and spatial neuronal responses as mice performed the task. (a) Six example CA1 pyramidal cells display varying extents of non-spatial (*left-most*) to spatial (*right-most cells*) tuning. *Top,* Trajectories (gray lines) separated by trial type. Only correct trials have been included here. Spikes from a single neuron are overlaid in *red*. *Bottom,* Average firing rates are shown for each trial type. Each cell generated a firing field where the firing rate was calculated with respect to position, shown for NT and TONE trials separately. In addition, a “frequency map” was generated for each cell, where the firing rate was calculated in relation to the auditory frequency. Average rates across trials are shown as heatmaps. *Rainbow spectrum*, trial types for ports 1-6 during TONE trials. (b) All cells with a significant firing field in the frequency map (for frequency field definition, see Experimental Procedures) are ordered by frequency preference. Many cells had non-spatial tuning (vertical stripe-like pattern in the position map generated by averaging across trials) and also showed faint stripe-like firing in the NT trials. While most tone cells responded near the target frequency, middle frequency-related responses were also observed (see *Cell 2*). Cells 1 to 3 from *a* are highlighted on the *right*. (c) All cells with a significant firing field during TONE trials in the place map (averaged across Ports 1-6) are ordered by spatial preference. Spatial tuning persisted but was significantly altered between NT1 and TONE trials (see **Extended Data Fig. 6-7**). Cells 4 to 6 from *a* are highlighted on the *right*. (d) Using a P-GAM (for details see Experimental Procedures and **Extended Data Fig. 3**), >60% of active neurons were classified as significantly modulated by both tone frequency and space, suggesting conjunctive spatial and non-spatial responses. *Colored dots*, individual sessions from each animal; *colored lines*, average proportion for each animal (IZ39-48). (e) The relative weights of each neuron for two example assemblies, detected using ICA. Place cells are labeled in *black* and tone cells in *pink*. Assemblies 1 and 2 are dominated by place and tone cells, respectively. (f) *Top*, Activation profile of all detected assemblies from an example session surrounding the first detected lick during TONE trials, sorted by the time of assembly peak activation. Note that the tone cell modulated assembly 2 is most active immediately before the lick. *Bottom,* PSTH of activation around the lick time specifically for assemblies 1 (*green*) and 2 (*blue*). (g) The sum of weights of all place cells (*black*) and tone cells (*pink*) in an assembly plotted against the peak time of activation of that assembly around the choice lick. Place cells contributed most to the assemblies early during a trial, but “tone cell” assemblies emerged preceding the lick (comparison between the sum of “tone cell” weights and place cell weights at each time bin, n = 33 sessions from 6 mice, Wilcoxon signed rank test. *Black lines* are when the weights of place assemblies were significantly greater than tone, *pink lines* are vice versa) *\*p<0.05,***p<0.001.* Line plots show mean ± SEM.

In addition to tone cells, there were clear examples of cells firing in fixed spatial coordinates regardless of the mouse’s choice and auditory tones (**Fig. 2a, c, Cells 4-6**). Upon closer inspection, these place cells were additionally modulated by the mouse’s approach to different ports during TONE trials. During TONE trials, several cells shifted or expanded their place fields with varying run lengths (**Extended Data Fig. 2h, i**). These results suggest that rather than belonging to unique functional cell classes, both tone cell and place cell neurons can have mixed spatial and non-spatial properties.

To quantify to what extent cells were comodulated by space and progression to the chosen port, we used a Poisson Generalized Additive Model (P-GAM;(Balzani et al., 2020; Noel et al., 2022)), which modeled the average response of a neuron as a function of continuous behavior variables such as spatial position during NT and TONE trials and the progression to the choice port (or tone frequency) (**Extended Data Fig. 3**). Almost 60% of the cells were co-tuned to both position and tone frequency (**Fig. 2d**). The ratio of the mutual information (MI) for position versus frequency revealed a continuous distribution (**Extended Data Fig. 3h, i)**, establishing that rather than existing as discrete functional classes, most cells exhibited tuning preferences lying along a continuum. Consistent with this idea, place cells and tone cells displayed prototypical properties of hippocampal cells, such as theta phase precession (**Extended Data Fig. 4**) and theta compression, with place and tone cells co-firing within the same theta cycle (**Extended Data Fig. 5**).

Given the mixed nature of neuronal responses, we identified assemblies of co-firing neurons that were not limited to functionally defined classes (see Experimental Procedures (Lopes-dos-Santos et al., 2013)). Across approaches to Ports 1-6, approaches to distinct ports were associated with similar assembly sequences, but the target warped their temporal dynamics (**Extended Data Fig. 6a, b**). Linking these assemblies back to functional types, we observed that assemblies active early in a trial contained mostly place cells, while those at the end mainly tone cells (**Fig. 2 e-g**).

In addition to modulation of hippocampal firing between different TONE trials, we also observed significant remapping from NT to TONE trials, but not during return runs (**Extended Data Fig. 6c-f**), indicating context-dependent hippocampal patterns. To address the source of remapping, we trained a separate cohort of mice (range of neurons: 59-194 cells; 2049 cells total with n = 1718 pyramidal cells and n = 331 putative interneurons out of 16 sessions in 3 mice) on a control task in which the reward was kept fixed to the end port even during TONE trials (**Extended Data Fig. 6g-i**). Therefore, auditory sweeps were identical but had no task-related significance. As expected, there was a time-proportional drift in neuronal firing throughout the session, yet no remapping was observed between NT1 and TONE trials (**Extended Data Fig. 6j-l**). Thus, sound alone could not explain the remapping.

Mice ran stereotyped trajectories during NT trials, mainly along the wall of the track without water ports. In contrast, during TONE trials and the second block of NT trials, the running trajectories were consistently closer to the wall with middle ports, reflective of the mouse’s search for the correct ports **(Extended Data Fig. 7a, b**). We regarded the linear track as two-dimensional space and examined the impact of minor deviations (Liberti et al., 2022). A decorrelation was observed between NT1 and NT2 trials in the two-dimensional maps (**Extended Data Fig. 7c-e**). We hypothesized that a sequence of positions (where the mice are coming from or are planning to go) was responsible for this difference. We used hierarchical clustering (see Methods,(Liberti et al., 2022)) to classify the movement trajectories in the forward running direction into two groups: away from middle water ports (ports 2-5; **Extended Data Fig. 7f,** *gray lines*), or closer to the middle water ports (**Extended Data Fig. 7f,** *black lines*). Rate maps, averaging trials for each movement trajectory group and condition (TONE vs NT), were separately generated and correlated. Different movement trajectories yielded significantly different firing patterns within the same task conditions **(Extended Data Fig. 7g, h**) as expected under the spatial map framework. However, even the same trajectories but in different conditions led to remapping (**Extended Data Fig. 7g, h**), pointing to the impact of task engagement to remapping.

### External variables and their direct influence on hippocampal firing

Examination of error trials revealed that firing was not driven by auditory tones but rather by the animal’s choice. Nonetheless, it is possible that tone cell firing reflected the mouse’s “belief” or falsely perceived tones. We thus examined the activity of tone cells during probe trials, where the tone ramp was terminated at 4 kHz. Despite the truncated auditory cue, tone cells continued to ramp after the termination of the tone while the mouse approached the port (**Fig. 3a-c**). The spatial map, cell assemblies and neural sequences also remained stable despite the truncation of auditory cues **(Extended Data Fig. 8)**. Thus, neuronal firing was not induced and sustained by sound *per se*. Instead, the findings are more parsimoniously explained by an internally generated sequence toward an upcoming choice.

We observed no reliable relationship between firing rates and speed (**Extended Data Fig. 9j-l;**(McClain et al., 2019)), suggesting that tone cells were not speed-modulated. Because most tone cells fired immediately before approaching a water port, we next considered whether they were directly responding to the presence of the reward (Gauthier and Tank, 2018; Sosa et al., 2023). However, we found no difference between correct (*water reward delivered*) and incorrect trials (*no water reward delivered*) up to the first detected lick (**Extended Data Fig. 9a**). To further explore the extent to which rewards, or their expectation, might impact firing, we compared firing patterns on return runs. Throughout training, mice were rewarded at the home port only if the trial was correct and unrewarded after incorrect choices. Despite the expectation of the reward presumably being different between correct and incorrect trials, we observed no differences in cell firing on the return runs (**Extended Data Fig. 9h, i**). To follow up on these results, we designed a new task. Mice were trained to run back and forth on a linear track with a reward port only at one end (666 cells total with n = 527 pyramidal cells and n = 139 putative interneurons out of 2 sessions in 2 mice), therefore never having received a reward at the other end. Midway through the session, rewards were provided at both ends of the track (**Fig. 3d**). No changes in hippocampal firing were observed upon the introduction of the reward (**Fig. 3 e-g, Extended Data Fig. 10**), suggesting a limited direct influence of rewards on hippocampal firing.

**Figure 3:**
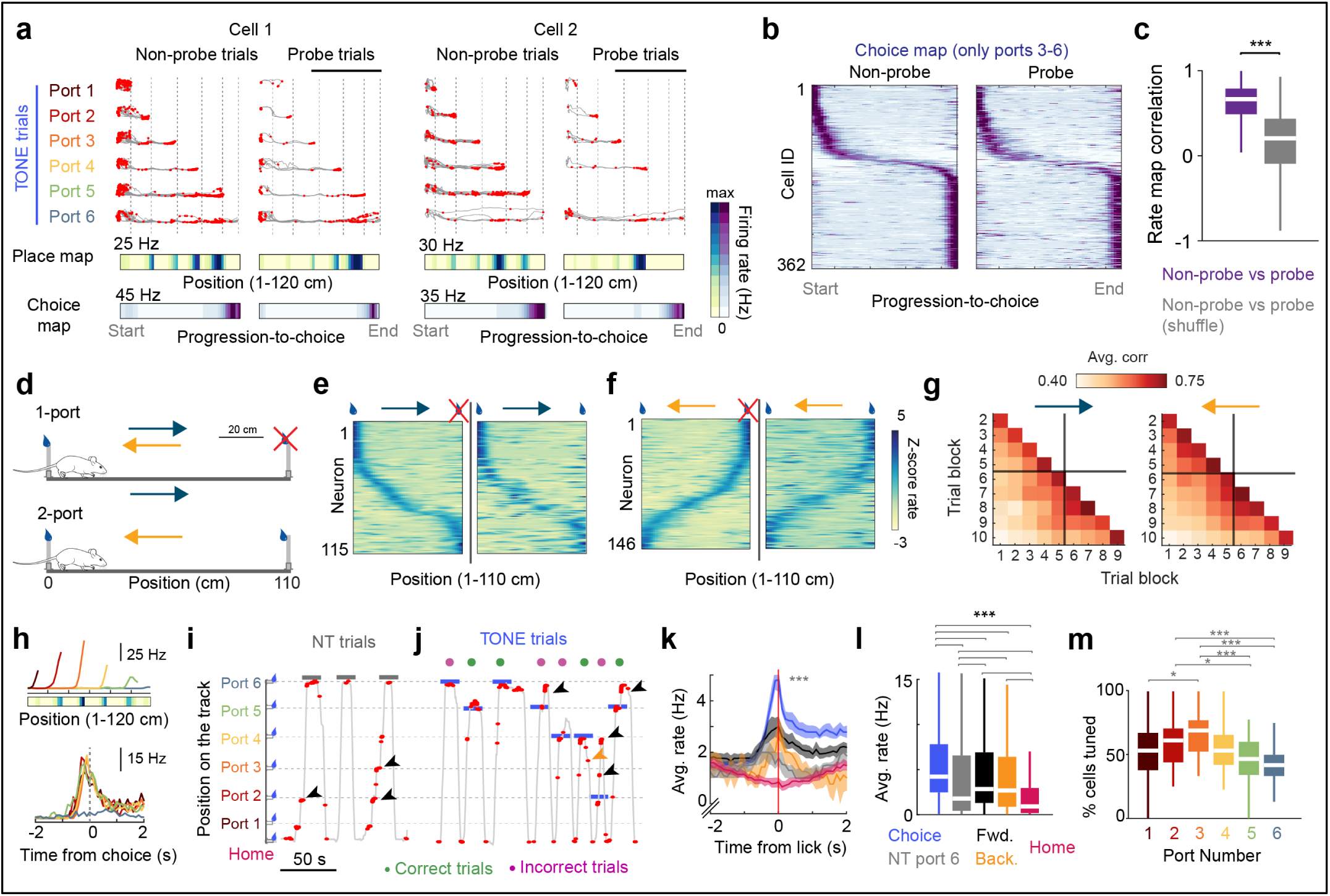
Hippocampal non-spatial firing is not driven by tones or rewards. (a) Two example CA1 pyramidal tone cells, recorded on sessions with probe trials (frequency truncated at 4 kHz). Probe trials were restricted to target ports 3-6 (*black horizontal line on top of the trajectory traces*) to ensure sampling of longer distances on the track after tones were truncated. *Left*, regular (non-probe) trials, *Right*, probe trials. Trajectories include correct and incorrect trials and are grouped by the chosen port for each trial. Note the similar distribution of the spikes at ports 3-6, even though the tone was already discontinued. Frequency maps in Fig. 2 are replaced by ‘choice maps’, calculated by scaling the trajectories to the mouse’s chosen port, reflecting the normalized distance to the choice (*progression-to-choice, 0-1 or start-end*). (b) Tone cells are sorted by tuning location during non-probe trials. Despite truncating tones, the maps remained stable between non-probe and probe trials. Thus, these cells did not respond to the auditory cue (also see **Extended Data Fig. 8**). (c) Cell-by-cell correlations for the cells in (b) compared to shuffled data (*gray*). Progression-to-choice activity remained stable despite the termination of the auditory tone (n = 362 cells from 8 sessions in 2 mice, Wilcoxon rank sum test against shuffled data, z = 23.23, p = 2.09 x 10-119). (d) Schematic of a control task to disambiguate the role of rewards on hippocampal firing. Mice ran 40-60 trials on a linear track where only one end was rewarded. Midway in the session, a reward was introduced on the non-rewarded end as well and the mice ran for another 40-60 trials. (e) *Left,* average rate maps of all place cells during the one-port condition (runs toward the variable reward port). *Right,* same cells in the two-port condition. Sorting is performed on the one-port condition and maintained between conditions. (f) Same as (e) but for place cells detected when the mice are running away from the variable reward port. (g) Average cell-by-cell correlation of rate maps for cells shown in (e, f) where maps were generated for 10 successive trial blocks. The reward was introduced at the beginning of block 6. Note a general decorrelation over time, but no significant decorrelation between blocks 5 and 6 for either running direction. (h) *Top,* Spatial rate map of an example “tone cell,” exemplifying the progression-to-choice tuning. *Bottom,* Peri-stimulus time histogram (PSTH) of the same cell with respect to the time from the lick from ports 1-6 (*rainbow colors* represent distinct ports as in *a*). (i) Spiking activity of the cell in (h) during NT trials. *Gray line*, spatial position of the mouse against time. *Red dots*, single spikes. *Black arrowheads* indicate times when the mouse spontaneously licked at the middle ports while running in the forward direction. Despite no tone present, the cell fired before each lick. *Gray lines* indicate licks at port 6. (j) Same as (i) but showing a segment of the TONE trials with correct (*green*) and incorrect (*pink*) trials. *Blue lines* indicate which port the mouse licked in each trial. *Black arrowheads* indicate spontaneous licks (i.e., sampling other ports after the mouse has already made a choice for that trial). *Yellow arrowheads* indicate spontaneous licks detected on the return run. (k) Average PSTH of reward-related tone cells, separated by the distinct lick types as shown in (i) and (j). Statistical comparison is between the average firing rate of the cells within ± 0.3 s around the lick. *Colors* as in (l). Lines shows median ± bootstrapped confidence intervals. (l) Average firing rate of the cells in (k) within ± 0.3 s around the first lick, separated by lick type (n = 565-633 cells from 6 mice, Kruskal Wallis test followed by Tukey-Kramer post-hoc tests, Chi-sq (4,3079) = 330.897, *p = 2.33 x 10^-70^*). (m) Fraction of cells per session that fired significantly before the approach to ports 1-6 (*rainbow shades*) during TONE choice licks. Later ports had fewer cells firing prior to the approach (n = 37 sessions from 6 mice, Friedman test followed by Tukey-Kramer post-hoc tests, Chi-sq (5, 180) = 35.44, p = 1.23 x 10^-6^). *\*p<0.05, ***p<0.001.* All box plots show median ± interquartile; whiskers show range excluding outliers.

Finally, we asked whether firing patterns were modulated by the expectation of reward by separating the type of lick the mouse performed. We grouped licks into five classes (**Fig. 3h-m**): (1) ‘Choice licks,’ the first detected lick on a TONE trial, which included both error and correct licks. (2) ‘NT licks,’ the licks at port 6 during NT trials. (3) ’Spontaneous forward licks,’ mice would sometimes sample other ports during the NT trials or after choosing in TONE trials. These spontaneous licks had no associated tasks or tones and may have reflected exploration or validation. (4) ‘Spontaneous reverse licks,’ same as type (3) but associated with licks that occurred as the mouse ran in the reverse direction back to the home port, meant to control for running direction and stereotyped motor movements. (5) ‘Homeport licks,’ the first lick detected at the home port that initiated the subsequent trial. Spontaneous (exploration) licks led to ramp firing similar to choice licks (**Fig. 3k, Extended Data Fig. 9d, e**). Strikingly, only weak responses were observed for NT licks and the smallest response was for homeport licks, suggesting this was not a uniform reward-expectation signal, and distinguished between habitual and task-modulated licks. During TONE trials, lick responses were not uniformly distributed across the approaches to all ports. Because the number of possible options sequentially decreased with every port crossed, the probability of a reward increased for later ports from the perspective of the animal. Licking at earlier (low probability) ports elicited strong responses, and progressively diminished for later (high probability) ports (**Fig. 3m, Extended Data Fig. 9f, g**). Neuronal firing, therefore, reflected a prediction signal for upcoming action outcomes, modulated by expectation, similar to the postulated role of dopamine(Bogacz, 2020; Mohebi et al., 2019; Syed et al., 2015).

**Figure 4:**
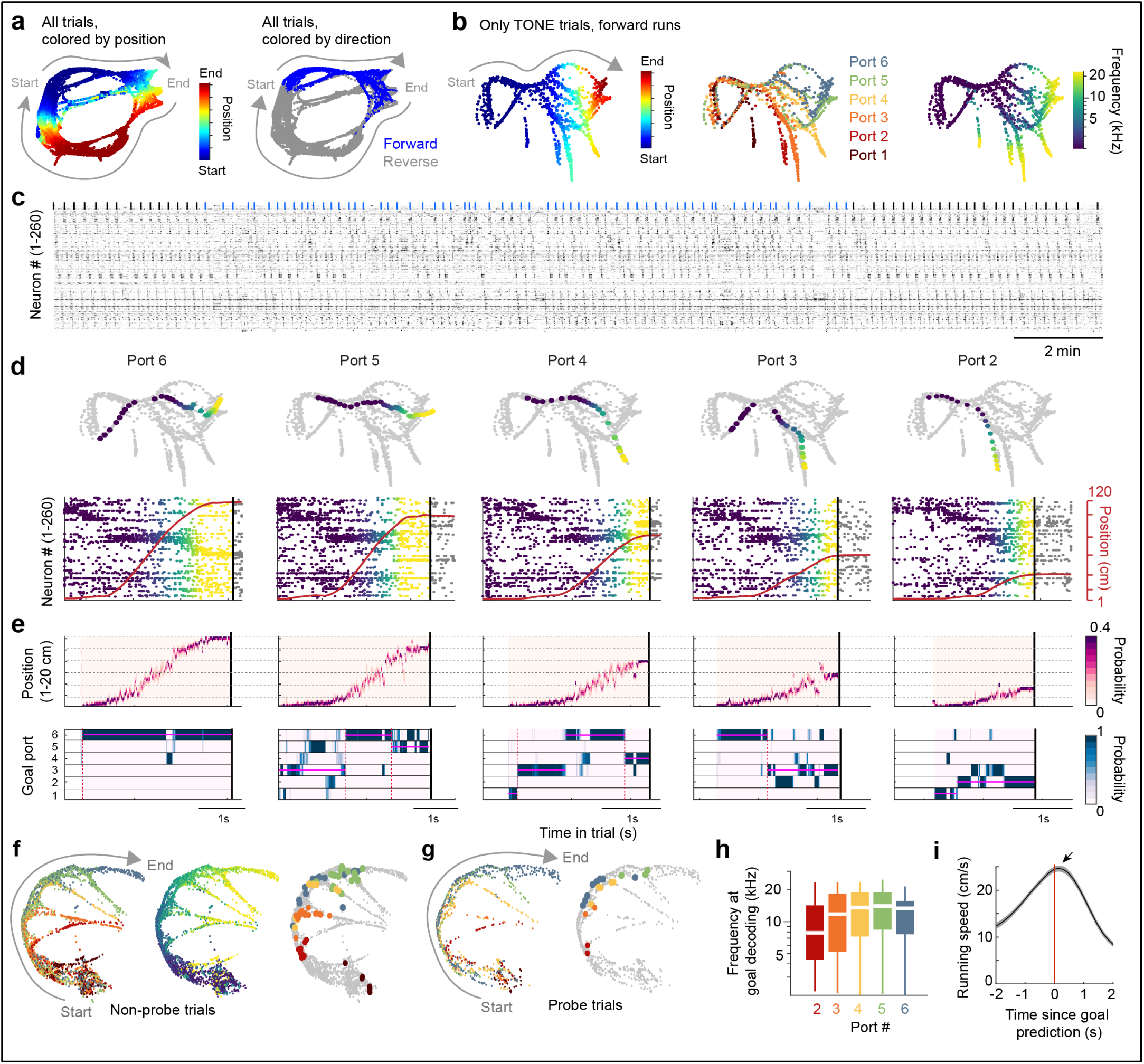
Hippocampal CA1 activity predicts upcoming choices. (a) Neural manifold from an example session. The activity of all neurons (n = 372 cells) was binned into 100 ms time bins and visualized on a neural manifold using UMAP. Each dot corresponds to a single time bin colored according to behavior variables. *Left*, Neural manifold color-labeled by the spatial position of the mouse on the track. *Right*, Same manifold, color-labeled by running direction. (b) Neural manifold from the same session as *a*, but only plotting the forward running directions during correct TONE trials until the first lick was detected. *Left*, spatial position; *Middle*, target port for that trial; *Right*, auditory frequency. Neural activity progressed along the main trajectory across trials but diverged to a unique path before the mouse chose a port to lick. (c) Spike raster of all active pyramidal cells from the session in (a), sorted by rastermap (see Experimental Procedures). Ticks indicate the start of NT (black) and TONE (blue) trials. (d) *Top*, UMAP (*gray dots*) as shown in (b), overlaid with a single trial going to the target port (colors correspond to auditory frequency; colormap in b). *Bottom*, spike rasters of the same trials, sorted as in (c) and colored according to frequency. *Vertical black line* indicates the time of lick, *red line* is the animal’s position. (e) 2-D spatial position and goal decoding (see Experimental Procedures) using 10 ms time bins. *Top*, Position decoding for the same trials as in (d). *Horizontal dashed lines*, port locations 1-6. *Bottom*, goal decoding for the same trials is in (d). *Vertical black line* indicates the time of lick for that trial. *Vertical dashed red line* indicates significant change points (see Experimental Procedures), when the goal decoding switches. *Pink horizontal lines* show the mode goal decoded between two change points. (f) UMAP as shown in (b), but for a different animal. *Left*, dots colored by the target port, colors as in (b). *Middle*, dots colored by frequency, colormap as in (b). *Right*, all dots shown in *gray*. Superimposed dots reflect the timestamp of the final detected change point during correct trials (see (e)). *Colors* indicate trials corresponding to ports 1-6. Decoding for each trial fell on the branch points for each target on the manifold. (g) UMAP from the same session as (f), only for 4 kHz probe trials. The UMAP shape remained similar, and goal decoding similarly overlapped with the branch points. (h) Average frequency at which the decoder settled onto its final change point to correctly predict the target, separated by port (n = 158-346 decoded trials). Port 1 was excluded because of short trial durations. The frequency at goal-decoding roughly corresponds to the frequency at which mice could predict the reward port as estimated from the probe trials (compare with Fig. 1h, **Extended Data Fig. 1**). Box plots show median ± interquartile. (i) Goal-decoding preceded running speed peak (arrow) by ∼0.2 s, after which mice decelerated and approached the reward port (arrow; n = 37 sessions from 6 mice). Line shows mean ± sem.

### Hippocampal population activity reflects behavioral goals

Examining the co-firing of neurons revealed that goal-predictive signals could emerge earlier than estimated from the peak firing rate of tone cells. To understand how the combined firing of neurons could be related to behavioral decisions, we examined hippocampal firing across the population. The activity of all recorded cells was binned into 100 ms time bins and visualized by the dimensionality reduction technique UMAP (Uniform manifold approximation and projection (McInnes et al., 2018)). Each population vector time bin on this lower dimension manifold was color-coded according to behavior variables, such as the spatial position or the direction of running (**Fig. 4a**). As expected, the manifold mirrored the geometry of the spatial environment(Rubin et al., 2019; Tang et al., 2023; Yang et al., 2023; Zhang et al., 2023) with unique segregations between forward and reverse runs. By selecting the correct forward runs during tone trials and coloring by spatial position, auditory tones, or target ports, we observed that the neural trajectories diverged from the main branch before the mice approached the target port (**Fig. 4b; Extended Data Figure 11).** We observed similar results across mice and sessions with varying numbers of simultaneously recorded neurons (**Extended Data Fig. 12-14**).

To examine how neural firing relates to the branches on the neural manifold, we sorted neurons using an unsupervised clustering analysis (see Experimental Procedures, Rastermap; **Fig. 4c**) and found reliable cell assembly sequences across trial types. To quantify how these assemblies relate to the manifold shape, we implemented a two-dimensional Bayesian decoder (see Experimental Procedures, (Brown et al., 1998; Johnson and Redish, 2007) to simultaneously predict the mouse’s position and chosen port (‘goal’) in 10 ms time bins. Each analysis method, including the UMAP display, cell sequences, and Bayesian decoding, showed a similar evolution of neural activity for port-specific runs, including error probe trials (**Fig. 4d, e, Extended Data Fig. 15**). Specifically, the mouse’s chosen port could be predicted for 85% of trials, ∼1.1 s before the lick (**Extended Data Fig. 16 a-d**). To relate this decoding time-point to branches on the neural manifold, we used a change-point detection algorithm ((Killick et al., 2012; Zheng et al., 2024), see Experimental Procedures) to identify consistent chunks of decoding prediction. The time point of the last-detected chunk generally occupied the branch points of the manifold for each port (**Fig. 4f, g, Extended Data Fig. 16e**). The average tone frequency at this time point was between 8-12 kHz (**Fig. 4h**), corresponding to the frequencies when the mice predicted the target port on probe trials (**Fig. 1h**). Importantly, decoding preceded changes in running speed (**Fig. 4i**). Thus, the goal-specific modulation of place and tone cells led to a population level code that corresponded to the animal’s upcoming actions.

### Within-trial evolution of action plan

We reasoned that if hippocampal activity reflected action plans, we could trace how such action plans are shaped within a trial (**Fig. 5a-d, Movie 3**). We identified moments of peak deceleration in the mouse’s speed profile when it approached the chosen port or crossed previous ports (**Fig. 5e, Extended Data Fig. 17a, b**). These transient deceleration events were often accompanied by brief head turns towards the port under consideration (‘deliberation’) (**Fig. 5b, Movie 3**). During deliberative but not during other decelerations on the track, tone cells briefly ramped their activity but quickly stopped spiking, corresponding to the mouse’s passing the port (**Fig. 5b, c, f),** which was also accompanied by a sudden jump in goal decoding (**Fig. 5b, c, g, Extended Data Fig. 17c**). While choice decelerations deeply extended into the manifold branches, deliberation neural trajectories only briefly entered the relevant manifold branch and exited back out (**Fig. 5d, h**). In addition, the likelihood of the animal’s position being decoded beyond the next port was significantly higher if the mouse was going to pass an upcoming port – an effect that was not dependent on the mouse’s speed but on its position ‘lookahead’ (Kay et al., 2020; Wikenheiser and Redish, 2015) during the ascending phase of theta cycle (**Fig. 5i, Extended Data Fig. 17d-g**). Thus, goal-specific signals were modified dynamically with ongoing action plans.

**Figure 5:**
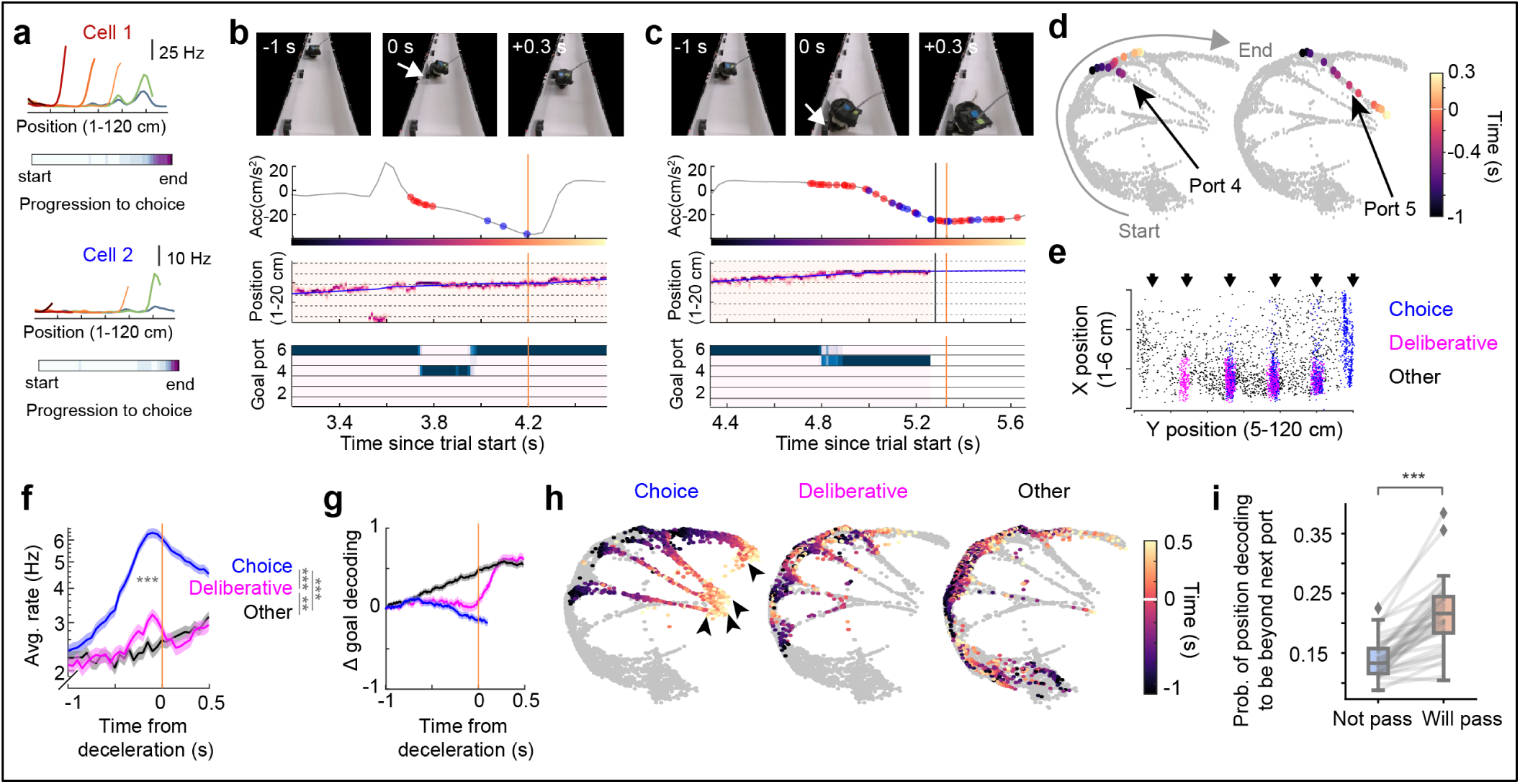
Action plan decoding in single trials. (a) Two example tone cells. *Top,* spatial firing for approaches to individual ports (rainbow shades). *Bottom,* Average progression-to-choice map. (b) and (c) Example events from a single trial, showing behaviorally detected “checking” of port 4 (white arrow in b; the mouse did not lick at the port) and licking at the correct port 5 (white arrow in c). (b) *Row 1*, frames from a front camera. *Row 2*, acceleration profile of the mouse (yellow line, acceleration minimum). Red and blue dots are spikes from cells 1 and 2 in (a). *Row 3,* position decoding for the same time window. *Blue line*, actual position of the mouse. *Row 4,* goal decoding for the same time window. (c) A second acceleration minimum occurred before the mouse approached and licked at the correct port 5. *Vertical black line*, time of choice lick. (see Movie 3 for a complete video) (c) UMAP display for the same session as Fig. 4f and the single trial shown in (b) and (c). Colored dots reflect points on the manifold corresponding to the time window shown in (*b, left*) and *(c, right).* Before port 4, the population deceleration only transiently entered the branch (*left*), whereas choice deceleration occurred well into the branch of port 5 (*right*). (d) For all mice, significant decelerations during the forward runs were detected (see Experimental Procedures) and clustered into three categories depending on where they occurred. Choice decelerations (*blue*), occurred upon the approach to a chosen port for a trial. “Deliberative” decelerations (*pink*) occurred near port locations (*black arrows*), but without licking. All other decelerations were labelled as ‘Other’ (*black*). Only ports 3-6 are considered to allow for deliberation at previous ports. (e) Average tone cell firing around each of the detected deceleration moments. As expected, cells ramped their firing in the 1s before the choice deceleration (as in Fig. 3). However, a small peak was also observed for “deliberative” decelerations (*pink line*), which was rapidly corrected after the mouse continued without licking at that port. This was not observed for other decelerations outside of port zones (n = 633 cells, comparison between firing rates between -100 and 0 ms. Friedman test followed by Tukey-Kramer posthoc tests, Chi-sq (2,1264) = 399.23, *p = 2.04 x 10^-^*^87^). (g) Change in goal decoding over time compared to the predicted goal at -1s. “Deliberative” decelerations were accompanied by a sharp jump in goal decoding, suggesting a sudden shift in goal prediction at the population level. (h) Significant decelerations (same session as d), separated by category. *Left,* Choice decelerations reach the end of the branches (*arrows*), *Middle*, “deliberative” decelerations show jumps in and out of the branch manifolds. *Right,* other decelerations rarely enter the branches. (i) The probability of the position decoder to predict a location beyond the upcoming port was significantly higher if the mouse eventually crossed the port (will pass) versus if it licked at the port (not pass). This reflects a “lookahead” beyond the current location based on the animal’s target port (Wilcoxon signed rank test, p = 2.91 x 10^-11^). Lines are individual sessions (37 sessions from 6 mice). *\*p<0.05, **p<0.01, ***p<0.001.* Line graphs show mean ± sem.

## DISCUSSION

We studied how hippocampal neurons vary their apparent tuning to various modalities in a paradigm where space, auditory tones, rewards, and contexts were juxtaposed. We observed correlations between firing patterns versus spatial position, time, auditory modalities, reward, and context. Yet, in agreement with recent experiments that attempted to identify modality-specific or abstract space responses (Allen et al., 2016; Aronov et al., 2017; Breathnach, 1980; Fischler-Ruiz et al., 2021; Gauthier and Tank, 2018; Josselyn and Tonegawa, 2020; MacDonald et al., 2011; Manns et al., 2007; Nieh et al., 2021; Purandare and Mehta, 2023; Radvansky et al., 2021; Sun et al., 2020; Tuncdemir et al., 2022), we found that the dominance of particular correlations depends on task demands and, therefore, varies across experiments. Analysis of the animal’s movement paths uncovered that hippocampal firing was linked to momentary action patterns and was modulated by goal expectation rather than external variables. We hypothesize that apparent hippocampal tuning to abstract cognitive variables can be grounded in a framework where neuronal firing reflects an internally generated action plan afforded by the animal’s environmental conditions.

Although adjusting the target frequency of the discriminative sound by the animal was a critical task variable, our analysis revealed that it was not the “sound landscape” (Aronov et al., 2017) that brought about the firing changes of hippocampal neurons but, instead, the specific actions taken by the animal. First, tone-responding neurons increased their discharge at sampled water-port locations during NT trials. Second, during error trials, the neuron’s firing pattern reflected the mouse’s progression toward the erroneously chosen port rather than the auditory frequencies. Third, when the tone ramp was terminated at 4 to 16 kHz on probe trials, neuronal firing continued to ramp similarly as the mouse approached the port. The weak or no effect of sound on hippocampal neurons may be related to the observation that self-initiated sounds produce minimal responses even in the auditory system (Audette et al., 2022). Finally, the firing patterns of hippocampal neurons “remapped” (Leutgeb et al., 2005; Muller and Kubie, 1987) across TONE and NT trials of the main task. Yet, the presence of the same auditory sweep with no consequence on securing the reward had no impact on neuronal firing in control experiments. We hypothesize that the role of the discriminative ‘go’ signal is to select an action sequence with an associated neuronal trajectory, which proceeds according to internally organized motives, possibly organized over learning, rather than continuously controlled by sensory cues.

An alternative explanation of our findings is that neuronal trajectories were influenced and scaled by the presence of the reward and its expectation (Gauthier and Tank, 2018; Krishnan et al., 2022; Lee et al., 2020; Sosa et al., 2023). Consistent with this idea, hippocampal neurons started to ramp up their firing rates before licks, independent of the animal’s location on the track or the presence of a tone. Yet, control experiments distinguished firing pattern correlates of reward and sensory cues from motor responses at both the single neuron and population levels. First, the ramping of lick-responding neurons was identical during rewarded and either erroneous or learned non-rewarded trials, demonstrating that reward expectation and consumption were not critical. Second, in the TONE task, neurons that robustly responded before licking also responded before exploratory licks but only weakly in the NT task despite involving the same motor patterns. Conversely, very few tone cells responded during home port licks, even though they resulted in the same volume of water as rewarded TONE or NT trials. In addition, the fraction of responding neurons and the magnitude of the responses was different between ports. Finally, tone cells fired even during hesitation or deliberation without licks. Thus, neither the reward *pe se* nor the motor response of licking could explain hippocampal neuronal firing patterns.

Several observations support the action plan hypothesis. Sequences of cell assemblies consistently evolved ∼1s before the choice, including probe trials. Early in the trial, assemblies were dominated by “place cells” and gradually replaced by tone cells. While these labels approximate the experimenter-established relationships, they do not imply distinct mechanisms. We observed reliable goal decoding and branches on the manifold emerging at a similar timescale, also coinciding with the approximate frequency at which mice formed a decision. Together, our findings align better with an interpretation that the discriminative signal selected a neuronal sequence (or trajectory), which proceeded according to internally organized motives that guided the animal’s current action plan. This view is supported by the observation that the hippocampus continues to generate the same fractions of place cell sequences even in the absence of the grid cell-supporting medial entorhinal input (Hales et al., 2014; Miao et al., 2015; Ormond et al., 2015; Robinson et al., 2017; Zutshi et al., 2022). Building on previous research (Aronov et al., 2017; Radvansky et al., 2021), we hypothesize that external cues do not continuously drive neuronal spikes in the hippocampus (Zutshi et al., 2022) but rather serve to select and update preexisting (or attractor) trajectories appropriate to a given task phase.

The hypothesis of neuronal sequence-sustained internal goals is supported by previous observations demonstrating that the extent and forms of neuronal sequences in the hippocampus correlate with the direction and distances of the animal-set goals (Green et al., 2022; Lee et al., 2006; Nieh et al., 2021; Pastalkova et al., 2008; Radvansky et al., 2021; Sun et al., 2023; Wikenheiser and Redish, 2015). Our findings are also relevant to rescaling place cells with changes in the size of the environment (Diba and Buzsáki, 2008; Gothard et al., 1996; Wikenheiser and Redish, 2015) and the intrinsic switching of their fields between reference frames (Fenton and Muller, 1998; Jackson and Redish, 2007; Jezek et al., 2011; Kelemen and Fenton, 2016; Kentros et al., 2004; Markus et al., 1995; Muzzio et al., 2009; Radvansky et al., 2021; Rowland and Kentros, 2008), as well as to goal-predicting “splitter” cells and prediction in general (Frank et al., 2000; Stachenfeld et al., 2017; Wikenheiser and Redish, 2015; Wood et al., 2000). Each of these independently studied features may reflect the same underlying physiological mechanism if hippocampal firing is viewed from the lens of a dynamical sequence generator that progresses toward a task-related goal (Buzsáki and Tingley, 2018; El-Gaby et al., 2023; Rueckemann et al., 2021; Villette et al., 2015; Whittington et al., 2020). While our results primarily pertain to non-spatial aspects of firing, we hypothesize that similar action sequences may also underlie spatial firing that is sensitive to precise trajectories and the animal’s goals (Jackson and Redish, 2007; Liberti et al., 2022; Markus et al., 1995). In our experiments, the evolving internal goals were reflected by the quantal jumps of the population activity from port to port and associated behavioral deliberations (brief head movements) of the animal (Movie 3).

The concept of a goal expands beyond a mere physical location and encompasses any inherent endpoint that leads to a desired outcome, as implied by experiments in humans (Barron et al., 2013; Schacter and Addis, 2009; Viard et al., 2011;). Despite the vagueness of these terms, the distinction between reward and goal is subtle yet crucial, shifting the focus from an external stimulus or location to an internally driven, intentional, and deliberative target (Hok et al., 2007; Nyberg et al., 2022; Radvansky et al., 2021). Departing from the notion of ’representation’ of external stimuli or ’mapping’ the environment onto the hippocampus, we hypothesize that its internally generated dynamics are projected onto the animal’s niche to make sense of the world through its actions.

## METHODS

### Experimental Animals

The Institutional Animal Care and Use Committee at New York University Langone Medical Center approved all experiments. Mice were kept in the vivarium on a 12-hour light/ dark cycle and were housed 4-5 per cage before implantation. Upon the onset of behavior training and subsequent surgery, the mice were moved to a 12-hour reverse light cycle (lights on/off at 7 pm/am) and housed individually. Mice were provided food and water *ad libitum* but were water-restricted to maintain 80% of their weight during and after behavioral training. We used a combination of transgenic and wildtype male mice [Auditory task; n = 6 male mice, of which 4 were double transgenic mice crossed between Pvalb-IRES-Cre females (Jax Stock No. 017320) and Ai32 males (Jax Stock No. 024109). The other 2 mice were C57BL/6J wildtypes (Jax Stock No. 000664). Control task: n = 3 male mice of which 1 was double transgenic as described above, and 2 were wildtype]. No obvious differences in behavior or neural firing emerged between the two types of mice. At the time of implantation, mice ranged from ages of 3-6 months old and 22-31g in weight. Variable reward task: n = 2 male mice of which both were double transgenic as described above.

### Surgical Procedures

Mice were implanted with 128 channel silicon probes (Diagnostic Biochips ASSY-INT128) into CA1 (Task mice: 1 mouse was implanted with a P128-6 probe, while 5 mice were implanted with P64-1-D Janus double-sided probes; Control mice: 1 mouse was implanted with a P128-6 probe, 2 mice were implanted with P64-1-D Janus double-sided probes). For all surgical procedures, mice were induced with 2.5-3% isoflurane in oxygen anesthesia (SomnoSuite Low-Flow Anesthesia System) and placed in a stereotaxic apparatus (Kopf Instruments). The level of isoflurane was then decreased to 1.3-1.5% for the remaining surgical procedures. During the surgery, body temperature was maintained at 37°C using a heating pad (Physitemp, TCAT-2LV Animal temperature controller). Vaseline was applied as eye lubricant. Following no reaction to toe-pinch, a small incision was made in the scalp, the scalp was cleaned with iodine (Dynarex Povidine Iodine solution) and lidocaine cream (Ferndale LMX4) was locally applied. The skull was then scored using a scalpel to ensure adhesion to the headcap. The coordinates used for CA1 were -1.9mm AP, +1.6 mm ML from bregma. A ground screw coupled with a 0.005” stainless steel wire (A-M Systems, #792800) was implanted in the skull above the cerebellum. Next, a base plate was cemented to the scored skull surface using Metabond (Parkell C&B, #S380). The probes were mounted on custom-made metal micro-drives to allow precise adjustment of vertical position of sites after implantation(Vöröslakos et al., 2021). During surgery, probes were implanted at the level of the cortex (0.4 mm DV). A combination of mineral oil (Fisher chemical, #O121-1) and paraffin wax (Sigma-Aldrich, #18634) (melted in a 1:1 ratio) was used to seal the craniotomy and cover the probe shanks. In the transgenic mice, four 200 µm optic fibers were also implanted in the brain, but sessions when those stimulations were performed were not included in this manuscript. The microdrive was cemented on the metabond skull surface using a combination of light-cured and regular dental cement (Unifast LC, Unifast Trad). Finally, a custom 3-D printed grounded copper mesh hat was constructed(Vöröslakos et al., 2021), shielding the probes. Following surgery, an NSAID analgesic was injected (Ketoprofen at 5 mg/kg, ∼0.13ml for a 25g mouse, stock solution of 1 mg/ml, Subcutaneous). Mice were allowed to recover and were continuously monitored for at least a week before water restriction and behavior training resumed. Body weight was monitored for all the days following surgery.

The position of the probe was confirmed at the end of each experiment. The mice were perfused with cold saline solution (0.9%) followed by pre-made 4% paraformaldehyde (PFA) in phosphate-buffered saline (PBS) solution (Affymetrix USB). Brains were post-fixed for 24h in 4% PFA and sectioned coronally to visualize the hippocampus. Sections were obtained with 40-50 µm thickness using a vibrating blade microtome (Leica, VT1000S), mounted on electrostatic slides, coverslipped with DAPI Fluoromount-G (Southern Biotech, 0100-20), and imaged using a virtual slide microscope (Olympus, VS120). No additional immunohistochemistry or tissue processing was done to visualize the probe tracks.

### Behavioral apparatus and task description

We constructed a linear track using white acrylic (1/8” thick) with dimensions of 120 cm (length) x 7.5 cm (width), with 5 cm high walls. The track contained 7 equally spaced (20 cm apart) water ports. The home port and port 6 were located on the short walls of the track, while ports 1-5 were located along one of the long walls of the track. Water was delivered through blunt 18G needles connected (using Tygon E-3603 Tubing; 1/16” ID, 1/8” OD) to solenoid valves (Cole-Parmer, EW-98302-02) that were briefly opened for 30ms. The needle was positioned precisely within a U-shaped infrared (IR) sensor (HiLetgo LM393 Correlation Photoelectric Sensor), such that any licks on the needle would break the IR beam. Successive breaks at the same port during a visit were disregarded during analysis. A speaker (MakerHawk Mini Speaker, 3W 8Ω) was positioned above Port 6, facing the home port, and was calibrated to ensure that it played frequencies up to 25 kHz (60-80 dB SPL; 20dB roll-off between 2kHz and 25kHz).

Water delivery and the tones were controlled by a combination of Bonsai (https://bonsai-rx.org/) and a custom-made Arduino circuit. A Basler overhead camera (acA640-90gc, Graftek Imaging) acquired images with a frame rate of 30 Hz. Bonsai was used to detect the real-time position of the mouse by contrasting the mouse’s black fur against the white maze. This centroid position was sent to Arduino using the serialStringWrite function. The Arduino would randomly assign a target port for a trial and rescale the Bonsai-detected position of the mouse to the distance between the home and the target port. For example, if the target port is 6 (located 120 cm from the start location), and the mouse’s current position is at 60 cm, the instantaneous normalized position is 60/120, i.e., 0.5. However, if the target port is 4 (located 80 cm from the start location), the normalized position becomes 60/80, i.e., 0.75. Auditory frequencies were linked to this normalized position according to the following formula:

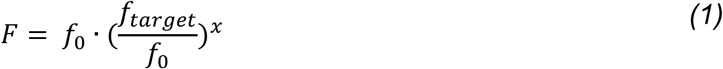

where, *F* = current frequency, *f*_0_ = 2 kHz, *f_target_* = 22 kHz, and *x*, is the current normalized position. The frequency sweep was designed to increase logarithmically from 2 kHz (similar to(Aronov et al., 2017)), with the target port coinciding with a frequency of 22 kHz. While tones could continue beyond the target, because of the limitations of the speaker, frequencies beyond 25 kHz were not accurately delivered if the mouse overshot, but these were rare instances. Tones were delivered using the Arduino Tone() function, which generated square pulses modulated at the specified frequency. Microphone (MAONO, AU-PM422, sampling rate up to 192kHz) recordings ensured that auditory frequencies were playing as designed and the resultant sound spectrogram was inspected using Audacity.

***Acoustic cue-guided navigation paradigm.*** During no-tone trials, mice ran along the linear track, receiving water at the two ends of the track, i.e., port 6 and the home port. In the tone trials, licking at the home port initiated a tone of 2 kHz that ascended logarithmically in a closed loop with the animal’s position. A correct lick response triggered the solenoid, and the tone stopped playing. We ensured that the delay between the mouse touching the spout and the TTL of the lick was not greater than 33 ms (corresponding to 1 frame-rate cycle of the Arduino) and corrected this delay in our analysis. After a choice, mice were free to sample other ports (*unrewarded checks*) before returning to the home port to lick to initiate the next trial. If the previous trial was correct, mice also received a water reward at the home port. However, if the previous trial was incorrect, both the choice lick and the subsequent home port lick were unrewarded. An incorrect lick response in the tone trials was followed by a 1-s long 3 kHz error tone.

***Probe trials***. In 2 mice, probe trials were randomly interspersed with regular (non-probe) trials at ∼30% probability, but only if the target was port 3, 4, 5, or 6. This criterion was introduced because ports 1-2 did not allow enough separation in terms of distance between the truncation of tones and the target port. Within a session, a cutoff frequency was pre-determined as 4kHz, 8 kHz, or 16 kHz, after which tones stopped playing with a trial. A correct probe trial was if the mouse correctly extrapolated the sweep speed and licked at the target based on the short, truncated sweep. Other licks were classified as incorrect. For electrophysiological recordings, only 4kHz probe trials were used to maximize the duration when sounds were off.

***Control task paradigm.*** To control for auditory cues, 3 mice were trained to run back and forth on the same track with the same task structure as described previously. However, only the two ports at either end were rewarded. Tones would stop playing after 25 kHz. Return runs did not have any tones associated with them. For comparison with the mice performing the task, recordings were only analyzed after mice had been performing the control version of the task for at least 7 days. This ensured that there were no effects due to the novelty of the auditory cue and that mice had learned that the tones were irrelevant.

***Variable reward paradigm.*** Mice had previously been trained to run in the auditory navigation task. After collecting data on that task, mice were kept in their home cage for 3-4 weeks. They were then introduced to a novel linear track of similar dimensions (110 cm long, 6 cm wide). Recordings were analyzed on the first day of this task. For the first ∼60 trials, only one end was rewarded. Midway through the task, and without interruptions, a second reward was introduced at the opposite end of the track.

### Behavioral training and analysis

On days when behavioral sessions were performed, the mice were water-deprived and only received water during the sessions. On other days, they were provided 2 ml of water. Body weight was monitored on all days, and the water schedule was interrupted if the body weight reached below 80% of the starting weight. Before implantation, mice were pre-trained to perform the task until they reached 65% accuracy. The stages of training included – (1) Habituation to the track and water ports, so mice learned to poke to trigger water rewards (∼2 days). (2) Linear track training where mice received a water reward at either end after poking and reached up to 60-70 trials (∼3-4 days). (3) Autoshaping, where tones would increase in a closed loop to the mouse’s position, but the water reward would automatically be released at the target port without the mice having to nose-poke. Incorrect licks were also not penalized (∼2 days). (4) Training to abstain from licking at incorrect ports. Only one port was assigned the target in this stage, generally port #3. Mice must poke at the target to receive water rewards, and licks at earlier ports would result in incorrect trials. This would often be the most variable stage between mice. If mice did not adapt their behavior after 100 trials, the first few ports would be physically blocked to train the mice to abstain from premature licks (∼3-4 days). (5) Expanding phase (4) to a few more ports, usually ports 1-3, or ports 3 and 6 (3-4 days). (6) Final version of the task with all ports rewarded (∼3-4 days). Mice transitioned between stages 4 to 6 if they reached 65% at each stage. The entire training from start to finish would usually take ∼3 weeks. Following surgery, mice were retrained starting from stage 4 but would quickly transition to stage 6 within 3-4 days.

To quantify the animals’ performance, the performance correct and proportion incorrect was calculated for each target port separately. The performance was calculated as the fraction of correct licks at a given port relative to the total number of trials where that was the target port. The proportion of incorrect trials was calculated as the fraction of trials when a given port was incorrectly chosen relative to all trials where that port was not the target.

### Electrophysiological recordings, unit clustering, and neuron classification

Electrophysiological data were acquired using an Intan RHD2000 system (Intan Technologies) digitized at 30 kHz. The wideband signal was downsampled to 1250 Hz and used as the LFP. A pulley system was designed to counteract the weight of the tether and headstage. The animal’s position was recorded using an overhead Basler camera (acA1300-60 gmNIR, Graftek Imaging) sampling at 30Hz. A front camera recorded the forward running direction of the animal for additional behavioral analysis (see Movie 1). Camera videos were synchronized with neural data with TTLs signaling shutter position. Digital inputs to the Intan RHD system provided timestamps for TTLs from lick ports and water delivery from the solenoids.

Spikes were extracted and classified using Kilosort (Pachitariu et al., 2016) using a custom pipeline KilosortWrapper (https://github.com/brendonw1/KilosortWrapper). Automated sorting was followed by manual curation of the waveform clusters using Phy (https://github.com/cortex-lab/phy) and custom plugins for Phy (https://github.com/petersenpeter/phy1-plugins). Kilosort clustering was performed with the following parameters: ops.Nfilt:6*numberChannels, ops.nt0:64;ops.whitening:’full’;ops.nSkipCov:1;ops.whiteningRange:64;ops.criterionNoiseChannels:0.00 001 ; ops.Nrank: 3; ops.nfullpasses: 6; ops.maxFR: 20000; ops.fshigh: 300; ops.ntbuff: 64; ops.scaleproc: 200; ops.Th: [4 10 10]; ops.lam: [5 20 20]; ops.nannealpasses: 4; ops.momentum: 1./[20 800]; ops.shuffle_clusters: 1. Units were separated into putative pyramidal cells and narrow waveform interneurons using their autocorrelograms, waveform characteristics and firing rate. This classification was performed using CellExplorer (Petersen and Buzsáki, 2020).

### Ratemap generation

1-D spatial rate maps were generated by binning spiking data into 2.5 cm wide bins, generating maps of spike counts and occupancy for periods when the animal’s speed was > 0.1 cm/s. A rate map was constructed by dividing the spike map by the occupancy map and smoothing it with a 5-bin Gaussian filter [0.02 0.1 0.16 0.1 0.02]. For TONE trials, forward direction maps used time windows between the start of the trial (home port lick), and end of the trial (first detected lick on the trial). Maps for the reverse direction runs included time windows between the end of a trial (first detected lick), and the subsequent home port lick. Because the mouse behavior was more variable during the return runs, an additional speed criterion was applied (speed < -2 cm/s) to ensure movement in the reverse direction.

1-D “frequency” or “choice” maps were constructed by rescaling the mouse’s spatial position to match the trajectory length within a trial, calculated as

(instantaneous spatial position/distance of the relevant port for that trial from the home port).

The relevant port was generally considered to be the port where the mouse first licked but was the assigned target port for the error trial analysis in Extended Data Fig. 2. For frequency maps, frequency was estimated using equation (1). Ratemaps were generated as described above but by binning spikes along the choice variable (bin size, 2.5% normalized distance to the choice port, also becomes a frequency map in the logarithmic scale). For TONE trials, average maps for ports 1-6 were first constructed separately, and then these average maps were averaged further to generate overall tuning across TONE trials.

2D spatial maps were generated by making 1-cm wide bins and binning across both the x, and y spatial positions. Smoothing involved a 5x5 bin Gaussian filter,

[0.0025 0.0125 0.0200 0.0125 0.0025;

0.0125 0.0625 0.1000 0.0625 0.0125;…

0.0200 0.1000 0.1600 0.1000 0.0200;…

0.0125 0.0625 0.1000 0.0625 0.0125;…

0.0025 0.0125 0.0200 0.0125 0.0025]

### Field detection

**<u>Place cells:</u>** Place fields were defined from the above-described rate maps using a combination of peak detection and across-trial correlation methods. Only pyramidal cells were used. The field boundaries were defined for cells with a minimum peak firing rate of 5 Hz, and extended till the rate was above 20% of the peak firing rate. The width of the field was required to be between 8% and 70% of the size of the ratemap (between 10 cm and 87.5 cm for place cells, and between 1.6 kHz and 14 kHz for tone cells). All cells that had a detected field using these criteria in spatial rate maps were included as place cells in **Fig. 2c**.

For the place field modulation by run length analysis shown in **Extended Data Fig. 2**, cells were only included if they had a place field in the ratemap generated by averaging across Port 2-6 TONE trials as well as a field within 30 cm of this averaged field for each individual trial type.

For the correlation metrics across NT and Port 6 TONE conditions, such as **Extended Data Fig. 6**, as well as the trajectory analysis in **Extended Data Fig. 7**, cells were included if they had a field in either NT or Port 6 TONE maps. Spatial rate-map correlations for individual cells between conditions were calculated as the average pixel-by pixel Pearson correlation coefficient of the smoothed-averaged firing rate maps. Rate map stability was defined by generating rate maps for the first and second halves of each condition and correlating these half-session rate maps.

The same analysis and field criterion were used for animals trained on the variable reward port task. For the correlation between the one-and two-port rewarded conditions, cells were included if they had a field in either map.

**<u>Tone cells:</u>** For “tone cells,” cells were selected with the same criteria as place cells to have a field in the averaged frequency maps. To ensure that these cells were firing robustly across trials, the frequency maps across ports 1-6 were correlated to each other and additionally required a correlation value >0.1. *Reward responses:* Peri-event time histograms around lick events were generated separately for each port for tone cells by binning spike rates in 100ms bins. To focus on cells firing around the reward location, only cells with a field within the last 10 bins were used for the analysis. *Speed analysis:* Speed analysis was performed by calculating the instantaneous firing rate of tone cells within 100 ms bins and calculating the Pearson’s correlation coefficient of the rate with running speed.

### Movement trajectory clustering

The movement trajectory of the mice along the width of the track was clustered similarly (Liberti et al., 2022) by using agglomerative hierarchical clustering. The x-position along the width of the track within a trial was downsampled to 7 points (each point corresponding to the y-position of a water port) for all NT and Port-6 TONE trials (total of n trials). This nx7 array representing individual x-position trajectories was clustered into paths according to their Euclidean distance using the *linkage* (Method, ‘complete’) and *cluster* functions in MATLAB. Only a maximum of 2 clusters was allowed. Two sets of average ratemaps were therefore generated for each condition (NT1, Port 6 TONE, NT2) by combining all trials classified within each cluster separately. We also generated within cluster, within condition maps by dividing trials by half, thus estimating baseline stability. These sets of ratemaps were then correlated within and across conditions.

### Poisson Generalized Additive Model

To estimate the tuning of each neuron to each variable, we used the Poisson Generalized Additive Model (P-GAM)(Balzani et al., 2020; Noel et al., 2022). The P-GAM defines a non-linear mapping between spike counts of a unit *y_t_* and a set of continuous variables *x_t_* and discrete events *Z_t_*. For our model, we selected continuous and discrete covariates - (1) the spatial position in the forward runs of the NT trials, (2) the spatial position in the forward runs of the TONE trials, (3) the spatial position in the return runs of all trials, (4) the progression to the chosen port (*progression-to-choice, or c*) during TONE trials, calculated as,

*p* = (instantaneous spatial position / distance of the chosen port for that trial from the home port)

To directly compare kernel strengths between (2) and (4), the *progression-to-choice* variable was re-scaled to the range of 0-120 cm(Singh and Singh, 2020). Finally, (5) The discrete covariate included the timestamps for the first lick for every port visit throughout the session.

The spike rates, continuous and discrete covariates were binned at a 33 ms resolution (30 Hz camera frame rate). Next, each tuning function was initialized as a basis set of cubic piecewise polynomials of order 4. The number of piecewise polynomials, or splines, was determined by the number of knots where the polynomials meet. The number of knots was chosen as k = 5 by an iterative process to uniformly cover the range of each variable without overfitting. See https://github.com/BalzaniEdoardo/PGAM for a tutorial.

The unit log-firing rate of each neuron was estimated as follows:

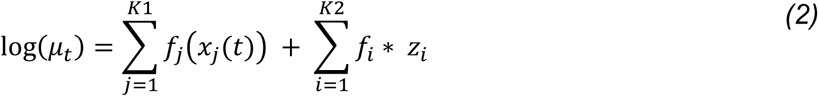

where *f_j_*(·) are non-linear functions of individual continuous input variables 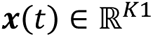 and discrete input variables 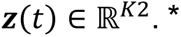. is the convolution operator. The spike rate of a neuron was estimated by a Poisson random process.

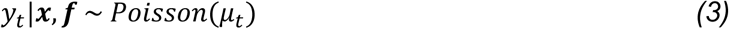

*f*(⋅) was modeled using a penalized spline basis expansion. We used flexible B-splines to define 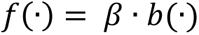, where b is the basis set as defined by the splines and *β* is a parameter that defines the weights or kernel strengths for each covariate. To enforce a smooth fit, each spline was associated with a quadratic penalty term ℒ(λ) to control its energy, or ‘wiggliness’. Both parameters β and the hyperparameters λ are learned from the data by an iterative optimization procedure (see (Balzani et al., 2020). To find the optimal regularization hyperparameter λ, a k-fold cross-validation (k = 5) was implemented before fitting the data using the Double Generalized Cross Validation score (dGCV). For each fold of the 5-fold cross-validation, the λ was selected for each neuron from the range of [1.5, 1.75, 2, 2.5, 5]. The final value of λ associated with the highest R^2^ was chosen to fit the model. The variability of the R^2^ values after each fold of the 5-fold cross-validation was visually inspected for 40 neurons of one session to ensure that the model fit was not sensitive to the subset of data being fit.

Our model was fit using 90% of the data, while neurons with a firing rate below 0.5 Hz were excluded from the fit. After fitting each variable, the model computed the marginal confidence intervals for the contribution of each variable to the neural response. The final values of the kernel strength and the magnitude of the confidence intervals were used to determine whether a variable has a significant contribution to the neuron’s firing rate. This was done using the Farebrothers algorithm, which computes the *p*-values associated with each variable. If the *p*-value was below the threshold of 0.001, then the variable was considered to contribute to the neural response significantly. After the initial fit of all variables (full model), the non-significant variables were removed from the model and the model was re-fit (reduced model), so that the parameters of only the significant variables were estimated again and more reliably.

In addition to the 5 variables described above, licks were also separated into 5 distinct classes as described in the text (choice, NT Port 6, Spontaneous forward and return, Home) and added as variables to the P-GAM fit. The tuning function of each lick class was initialized as a basis set of cubic piecewise polynomials of order 1 and 2 knots that were defined around the lick timestamp. The tuning function of the neuron to each lick class thus resembled a step function with a distinct *β*, or kernel strength for each type of lick, which allowed us to compare the modulation of the spike rate by lick type.

The Mutual Information (MI) was computed as the difference between the entropy of unit firing rates *H(Y)* and the entropy of the rate conditioned on a specific variable *X*, *H(Y|X),* under Poisson noise assumptions. Specifically,

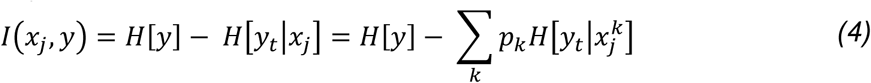

where *H*[*y*] is approximated as the entropy of a Poisson variable with λ equal to the mean firing rate of the unit. For *p_k_*(probability of the stimulus taking value 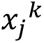), we used the empirical distribution of the discretized stimulus 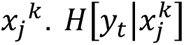 was approximated as the entropy of Poisson random variable with a mean given by the tuning function of the neuron, provided by the P-GAM model.

### Theta phase precession and theta compression

Average rate maps in the place and frequency domain were calculated for all TONE trials. Cells with a field in these averaged maps were then independently analyzed for fields within 30 cm of the averaged field in trials for ports 2-6 independently. LFP from str. oriens was filtered, and the instantaneous phase was determined using a Hilbert transform. Fields were normalized between 0 and 1 to the beginning to end of each field in either the space or tone domain. A circular-linear regression was generated for spikes occurring within the bounds of the fields, and the slope and significance of the regression line are reported. Significance was achieved by a p<0.05. Boundaries of ±2 cycles were used to constrain the circular-linear regression.

For theta compression, only averaged rate maps were used. The distance between the field peaks was estimated in either the space or tone domain for cells with overlapping fields. Spike times from pairs of neurons from within the field were cross-correlated with 5 ms bin size, and the peak lag between ±100 ms was estimated as the temporal lag between the cells.

### Assembly analysis

Cell assemblies were defined using an unsupervised statistical framework based on a hybrid PCA followed by ICA (Lopes-dos-Santos et al., 2013; van de Ven et al., 2016). Spike trains of pyramidal neurons were binned in 33-ms intervals, and the matrix of firing correlation coefficients for all pairs of neurons was constructed. Principal components with eigenvalues that exceeded the threshold for random firing correlations (using the Marčenko-Pastur law) were used to determine the number of assemblies. Next, using the fast-ICA algorithm (Ritchey et al., 2015), we determined the vector of weights (contribution of each neuron) for each assembly (component). Place cell and “tone cell” contributions for each assembly were estimated by averaging the weights of all place and tone cells for each assembly. The strength of each assembly’s activation as a function of time was determined by multiplying the convolved z-scored firing rate of a given neuron at a given time by the weight of that neuron’s contribution to the assembly. The product of these weighted spike counts was then summed for all nonidentical pairs of neurons to generate the instantaneous activation strength. Activations greater than 1.5 times the standard deviation were used as timestamps of activation of each assembly.

### Low dimensional manifold visualization with UMAP (unsupervised)

The procedure was implemented as in(Yang et al., 2023). Briefly, neural spiking data (spike count) during maze running (speed>1cm/s) were binned into 100 ms bins. The data were then smoothed using a 5 bin (500 ms) wide Gaussian kernel and z-scored. To examine the effect of binning and smoothing, we also performed finer binning (20 ms) and no smoothing (**Extended Data Fig. 11**). The UMAP dimensionality reduction algorithm was then applied to the preprocessed data matrix. Each point in the low-dimensional manifold corresponds to the population activity at a single time bin. The UMAP code is available at https://github.com/lmcinnes/umap/blob/master/doc/how_umap_works.rst

UMAP hyperparameters used were: n_neighbors = 20, metric = ’cosine’, output_metric = ‘euclidean’, learning_rate = 1.0, init = ‘spectral’, min_dist = 0.1, spread = 1.0, repulsion_strength = 1.0, negative_sample_rate = 5, target_metric = ‘categorical’, dens_lambda = 2.0, dens_frac = 0.3, dens_var_shift=0.1.

### Cell sequence visualization using rastermap

Neurons were sorted in **Fig. 4**, **Extended Data Fig. 8, and Extended Data Fig. 5** using rastermap (Stringer et al., 2023). Only pyramidal cells with an average firing rate >0.1 Hz were included. The same sorting was maintained for all figures. Specific parameters used are, n_clusters=50, n_PCs=200, locality=0, time_lag_window=3.

### Bayesian decoding

To decode the discretized position and chosen port of the animal at each 10ms time bin, we used a state space decoder that is the Bayesian decoder equipped with a causal filter on the posterior (Johnson and Redish, 2007; Johnson et al., 2008), based on specified transition probability on the position and goal (Brown et al., 1998). In the encoding stage, the spiking activity of the population is modeled as independent inhomogeneous Poisson processes whose instantaneous rates are functions (tuning curves) of the animal’s position and chosen port for the trial. The tuning curves (x) are constructed using a kernel density estimator with a Gaussian kernel of 3cm for the position dimension and virtually no smoothing for the goal dimension. The state transition probability *p*(*x_t+1_*|*x_t_*) is the product of the transition probability for each dimension, where the transition in the position follows a Gaussian random walk with a variance of 25cm^2^ per time bin, and the transition in goal follows a sticky uniform prior, where the probability of staying in the current goal is 0.9 per second (0.9∧0.01 per bin), and equal probability for the rest of the available goals (available meaning the goals that are ahead of the current position). The posterior of the state given the history of observations, follows this recursive relationship:

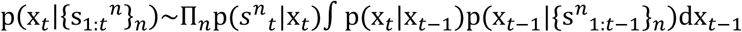

where the 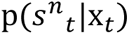 is the Poisson likelihood function of observing a spike count of 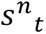 from neuron n at time t if the state variable is *x_t_*. The normalizer is the integral of the right-hand side with respect to *x_t_*, which is tractable because the state variables are discrete. Once we obtained the joint posterior of position and goal, we marginalized over one variable and analyzed the marginal posterior of the other. We adapted the code from the ‘replay_trajectory_classification’ Python library (Denovellis et al., 2021).

### Change point detection for goal decoding

To identify consistent chunks of goal decoding while ignoring minor fluctuations, we used the change point detection approach developed by (Killick et al., 2012) called pruned exact linear time (PELT) method (a modified version was used in (Zheng et al., 2024)). Briefly, the posterior for goal (marginalized over position) within a trial was partitioned into segments to minimize the sum of the within-segment variances plus a regularization term.

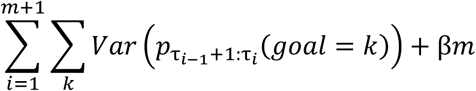

The regularization term is a penalty coefficient β x the number of change points *m*. The penalty coefficient was selected heuristically to produce large segments where the goal decoding was mostly stable (β = 20). Varying the coefficients tended not to change the starting time of the last segment, which was used to project onto the UMAP manifold. The changes points τ_1:m_ (τ_*m+1*_ defined to be the last time bin) that segment the posterior were found by dynamic programming with pruning (Killick et al., 2012). The number of change points was automatically selected by the tradeoff between the sum of variances and the regularization. The implementation used the Python library ‘ruptures’.

### Deceleration minima

During TONE trial forward runs, the mouse’s acceleration was convolved with a 200 ms Gaussian kernel. Using the MATLAB ‘findpeaks’ function, we identified troughs in the acceleration profile (‘MinPeakProminence’, 10). Each deceleration was then labeled as either the final ‘choice,’ deceleration close to a port before the final choice (‘deliberation’), or ‘other’ decelerations.

### Generalized Additive Model for theta lookahead

We use a generalized additive model (Hastie and Tibshirani, 1987) to predict the sum of the posterior beyond the upcoming port using linear covariates speed and will_pass (whether the animal will pass the upcoming port) and smooth terms position and theta phase.

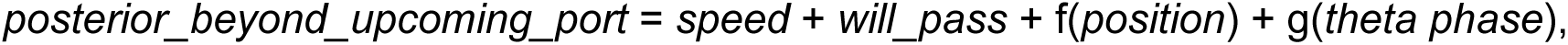

where, 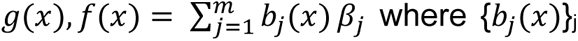 is a set of B-spline basis functions, and {*β_j_*}*j* are the coefficients. We used 30 basis functions for the *position* and 5 basis functions for *theta phase,* both with degree 3. The number of basis functions was heuristically chosen such that the fitted smooth functions were interpretable. Varying the numbers did not change the results regarding the significance of the *will_pass* term. We used an identity link function and Gaussian noise model. The model is implemented using the Python library statsmodels.

### Statistical analysis

Animals from the control paradigm and auditory task paradigm were run in parallel cohorts. No specific analysis was used to estimate a minimal population sample, but the number of animals, trials, and recorded cells were larger than or similar to those employed in previous studies (McKenzie et al., 2021; Senzai et al., 2019; Valero et al., 2021; Zhang et al., 2021). All statistical tests were conducted using MATLAB R2023b, and the details of the tests used are described with the results. Unless otherwise noted, all tests used non-parametric comparisons of means and variance (Wilcoxon paired signed rank tests, Wilcoxon rank-sum test, Kruskal-Wallis one-way analysis of variance, and Friedman tests). When parametric tests were used, the data satisfied the criteria for normality (Kolmogorov-Smirnov test) and equality of variance (Bartlett test for equal variance). All posthoc tests were performed using Tukey honest significant differences and correcting for multiple comparisons. Boxplots represent the median and 25^th^ and 75^th^ percentiles, and the whiskers represent the data range. In box plots without datapoints, outliers were excluded from the plots, but always included in the statistical analysis.

## DATA AVAILABILITY

All the data of this study will be made publicly available upon acceptance of the manuscript in the Buzsaki Lab Databank: https://buzsakilab.com/mp/public-data/.

## CODE AVAILABILITY

All custom data preprocessing code is freely available on the Buzsaki Lab repository: https://github.com/buzsakilab/buzcode. Code for the P-GAM implantation is freely available on https://github.com/BalzaniEdoardo/PGAM. Scripts specific to analyzing this dataset are available on I. Zutshi’s github page, https://github.com/IpshitaZutshi/JungleBook/tree/main/ToneTask.

## ACKNOWLEDGEMENTS

We thank R. Kasa and L. Anderson for help with behavioral training and Dr. A. Mar for help with the behavioral paradigm design. We also thank S. Sethi and the Buzsaki lab for insightful comments throughout the project and suggestions for the manuscript. Funding: This work has been supported by a Simons Collaboration on the Global Brain Transition to Independence Fellowship to I.Z., by the Bodossaki Foundation Scholarship for Postgraduate Studies and the Swiss-European Mobility Program for Worldwide Projects and Traineeships to A.A., an NIH NINDS grant 1RF1NS12712201 to E.B. and C.S., NIH grants (R01MH122391; U19NS107616) and an NSF grant 1707316 (NeuroNex MINT) to G.B.

## AUTHOR CONTRIBUTIONS

I.Z. and G.B. planned and designed the experiments. Experiments were performed by I.Z., A.A and T.D. Data analysis for single cells was performed by I.Z. and A.A. W.Y. performed the UMAP analysis. S.Z. performed the position and goal decoding analysis with support from A.W. E.B. and C.S. provided support for the P-GAM implementation. I.Z and G.B. wrote the paper with input from all authors.

---

**Extended Data Figure 1.**
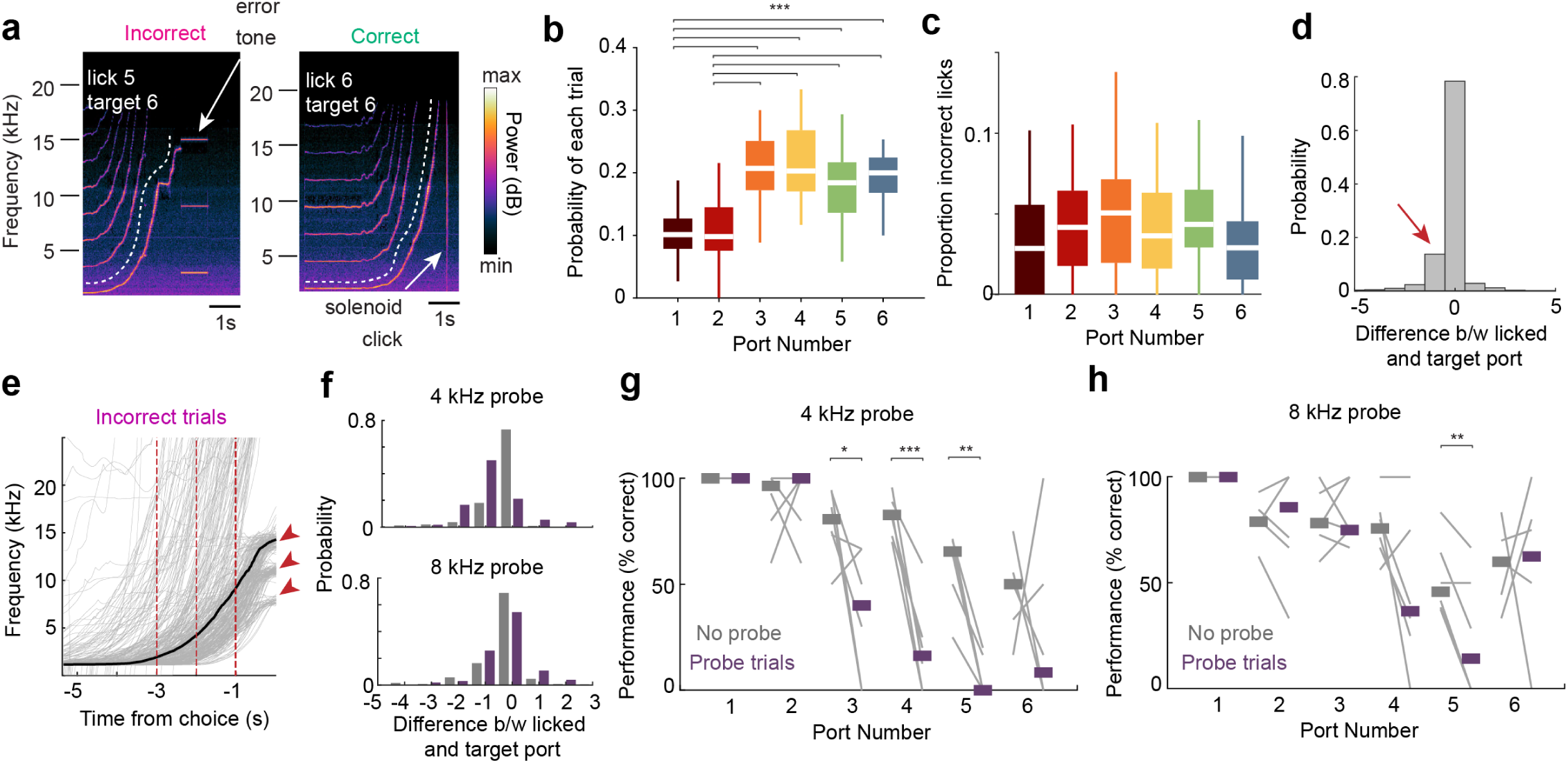
Behavioral variability in the task for ports 1-6. (a) Spectrogram of frequencies recorded in an incorrect and correct trial. Power below the *white dashed lines* represents the tones playing, while the power bands above the white lines correspond to analytical harmonics. A 3 kHz tone was played for 1s after mice made an erroneous lick (*left;* error tone), with no water delivery. Mice received a water reward if they poked at the correct port (*right*, white arrow, solenoid delivery made a clicking sound). (b) The probability of trials was deliberately unevenly distributed to yield more trials where mice performed longer runs on the track. Thus, the first two ports were less frequently rewarded than the later ones (Friedman test followed by Tukey-Kramer post-hoc tests, Chi-sq (5,180) = 61.47, *p = 6.02 x 10^-12^*). (c) The fraction of times the mouse incorrectly poked at a specific port compared to all trials where that port was not the target (n = 37 sessions from 6 mice, Friedman test followed by Tukey-Kramer post-hoc tests, Chi-sq (5, 180) = 9.50, *p = 0.0906*). (d) Histogram of the difference between the target and choice port. Most errors occurred when mice licked one port sooner than the target (*red arrow*). (e) Tone frequency progression of each trial (*gray lines*) in time, superimposed with the average frequency (*black*) for incorrect trials across all mice and sessions. If the mice overshot the target, the tone was cut off beyond 25 kHz. The discrete branches (*red arrowheads*) reflect errors one (top), two (middle), or three ports (bottom) before the target. *Red lines* are at -3, -2, and -1s before the first lick. Compare with Figure 1f. (f) Histogram of the difference between the choice and the target port for regular (“no probe*”; gray)* and probe *(purple)* trials. During 4 kHz trials, the mice generally made errors by licking earlier (“undershooting”), while during 8 kHz trials, the mice made both undershooting and overshooting errors. (g) Behavior performance during 4 kHz probe trials for each port. Impairment was only observed for ports 3-5. For port 6, the effect was variable, possibly because mice would interpret the truncation of the tone to signal a NT trial and run straight to port 6 (Paired t-test, n = 6 sessions from 2 mice, Port 1: *p ≈ 1;* Port 2: *p = 0.92,* Port 3: *p = 0.044;* Port 4: *p = 8.38 x 10^-4^*, Port 5: *p = 0.0012;* Port 6: *p = 0.*67). (h) Same as (g) but for 8 kHz probe trials. An impairment was observed for both Port 4 and Port 5, but only Port 5 reached significance (Paired t-test, n = 6 sessions from 2 mice, Port 1: *p ≈ 1;* Port 2: *p = 0.94,* Port 3: *p = 0.82;* Port 4: *p = 0.09*, Port 5: *p = 0.0084;* Port 6: *p = 0.*76). *\*p<0.05, **p<0.01, ***p<0.001.* All box plots show median ± interquartile; whiskers show range excluding outliers. Lines in g-h show median.

**Extended Data Figure 2.**
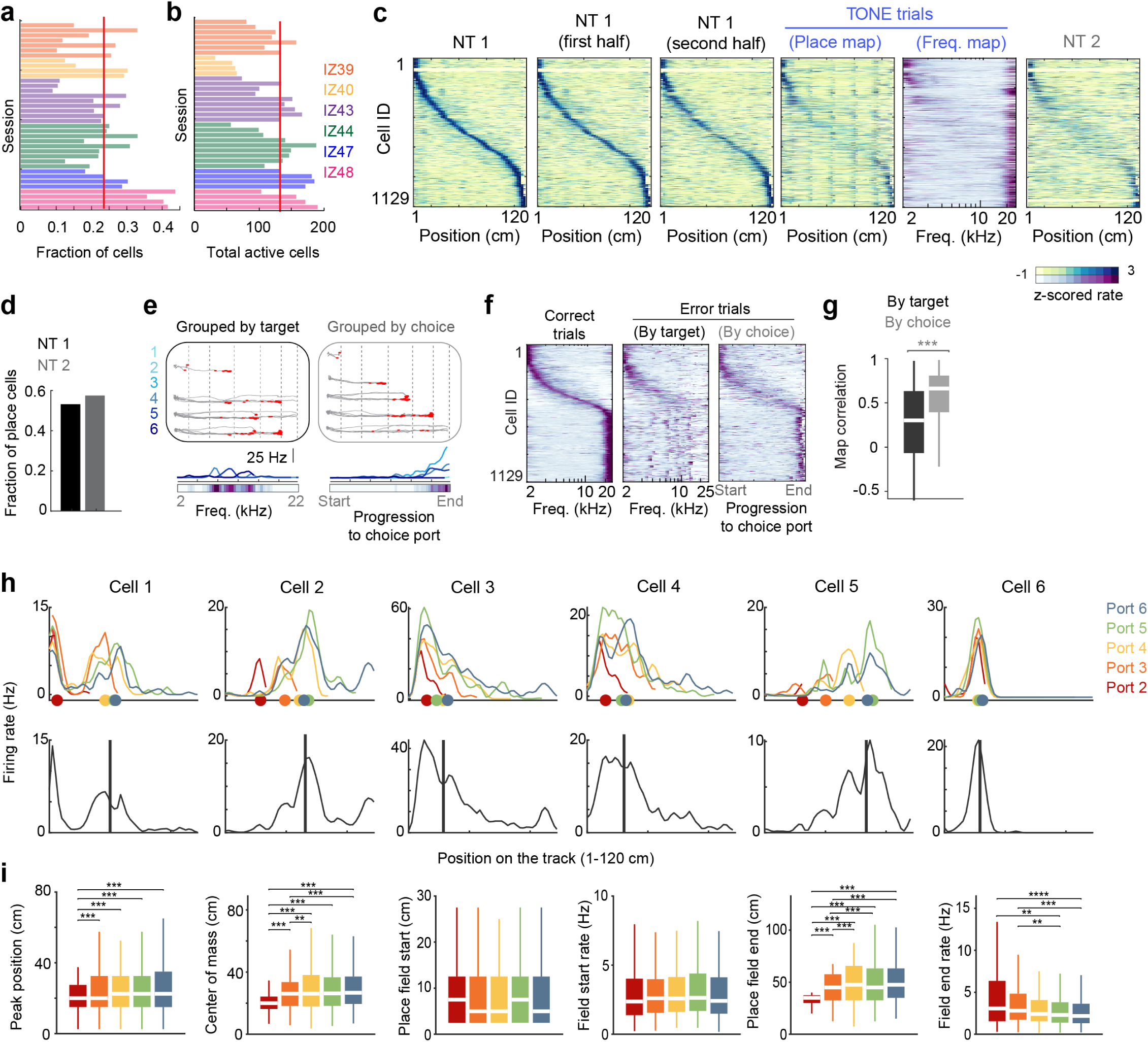
Modulation of hippocampal cells by space and approach to ports. (a) Fraction of pyramidal cells responding to the “tone” varied for each session (*bars*) and animal (*colors as in*). *Red line*, median. The successive color bars in each mouse correspond to subsequent sessions (top, earliest recorded session). (b) Total number of active pyramidal cells (average firing rate > 0.2 Hz) recorded in each session of (a). *Red line,* median. The successive color bars in each mouse correspond to subsequent sessions (top, earliest recorded session). (c) Same “tone cell” population as in **Fig. 2b**, but sorted according to spatial position preference during NT1 trials. Dividing NT1 trials into halves revealed stable place fields during NT1, but mainly non-spatial firing during TONE trials. Thus, depending on the context, tone cells could be spatially tuned on the same track. Sorting was maintained across display conditions. (d) A substantial fraction of tone cells had a definable place field in the NT trials, suggesting that tone cells are not a unique class. (e) Example firing of a tone cell on error trials. The average rate maps were generated by grouping trials by the target port or the choice port where the mouse incorrectly licked. Thus, these average maps either corresponded to auditory frequency (*left,* tuning to sensory variable) or the progression to the choice port (*right,* tuning to movement trajectory). The average ‘choice maps’ were calculated by scaling the trajectories to the mouse’s chosen port, reflecting the normalized distance to the choice (*progression-to-choice, 0-1 or start-end*). (f) The average firing of tone cells in error trials sorted as during correct trials. Averaging by the choice of the mouse, but not the target, more closely matched the firing of the same cells during correct trials. Thus, sound-induced firing reflects the movement trajectory to the choice port rather than responding to sensory variables. Colormap as in (c). (g) Cell-by-cell correlation of tone cells with their frequency maps during correct trials as in (b), versus during error trials, grouped by target (*e, left*) or choice (*e, right*). Colors as shown in (f) (n = 1129 cells from 37 sessions in 6 mice, Wilcoxon paired signed rank test, z = -18.96, *p = 3.47 x 10^-80^*). (h) Six example cells with a defined place field across all tone trials. *Bottom*, average rate map across all trials, *vertical black line*, center of mass of rate map. *Top*, colored lines, rate maps of the cell after separating trials by approach to ports 1-6. Colored dots, center of mass of each rate map. A variety of effects were observed, ranging from shifting place fields (Cell 1, 2), expanding place field tails (Cells 3, 4), truncating place fields (Cell 5), and stable fields (Cell 6). (i) Group statistics comparing port-specific rate maps of place cells. To ensure a fair comparison across all ports, cells must have a stable field in each port-specific rate map (Ports 2-6), which must be within 30 cm of the averaged field (*black line in h*). There was a significant effect of place field peak and the center of mass shifting (e.g., Cell 1, 2), place field tails expanding (e.g., Cell 3, 4), and place field rates at the end of the field decreasing, suggesting truncation (e.g., Cell 5). (Friedman test followed by Tukey-Kramer post-hoc tests, n = 321 cells, peak position: Chi-sq (4,1280) = 60.66, *p = 2.1 x 10^-12^*; center of mass: Chi-sq (4,1280) = 199.46 *p = 4.91 x 10^-42^;* start of place field: Chi-sq (4,1280) = 2.47, *p = 0.65*; place field start rate: Chi-sq (4,1280) = 7.91, *p = 0.095;* end of place field: Chi-sq (4,1280) = 353.68, *p = 2.81 x 10^-75^*; place field end rate: Chi-sq (4,1280) = 34.23, *p = 6.69 x 10^-7^*). *\*\*p<0.001, ***p<0.001.* All box plots show median ± interquartile; whiskers show range excluding outliers.

**Extended Data Figure 3.**
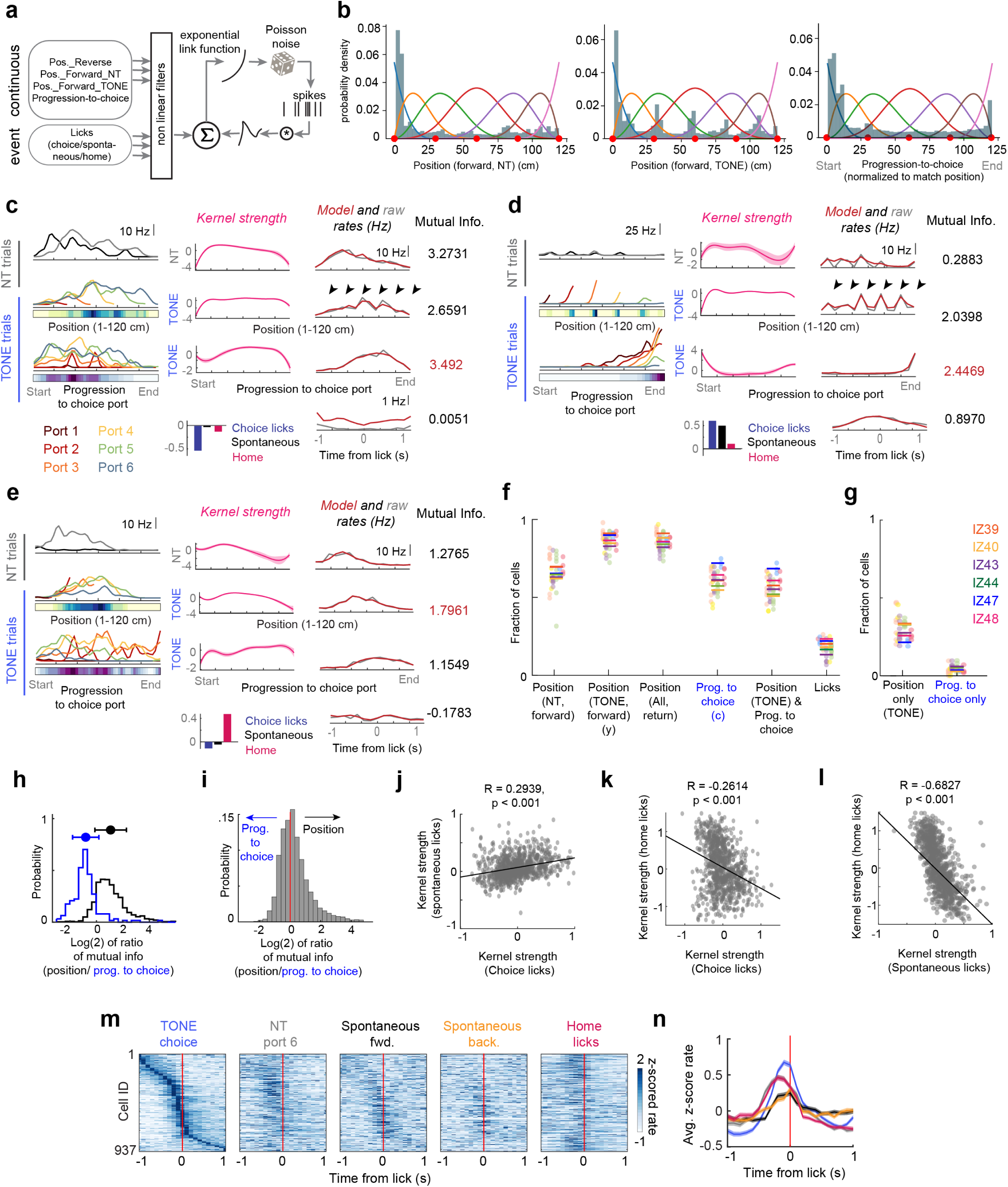
Fitting multiple behavior variables to model single-cell firing. (a) Schematic of the Poisson generalized additive model (P-GAM) used to fit spike trains. Inputs to the model included continuous task variables such as the position in NT and TONE trials for the forward and reverse runs and the progression-to-choice for the forward TONE trials. Event variables included all detected lick epochs, i.e., the first lick at a port visit. Subsequent licks within a visit were not included. Licks were grouped into choice during TONE trials, port 6 during NT trials, spontaneous and home port licks (see **Fig. 3 i, j**), and a step kernel function was fit for each lick type (see Methods). (b) Distribution of the continuous input variables for an example session. Knots (*red circles*) were defined to sample the distribution equally and B-splines (*colored lines*) for the P-GAM were constructed based on these knots. (c-e) Three example neurons modeled using the P-GAM. *Left,* average rate maps constructed from the spike trains for each continuous variable. *Middle*, Kernel strength (*β*), that, when convolved with the B-splines, generated the best fit of the neuron’s tuning curve. The kernel strength for the step function of each type of licks is shown as bars on the bottom plot. *Right*, True tuning curve (gray), and the tuning curve generated by the reduced model (*red*; i.e., only those variables that were significantly contributing to the firing of the neuron). Right, mutual information (MI) between the firing of the cell and each continuous variable. The highest MI for each cell is shown in *red*. The fit for licks was non-significant for the neuron in (c), but all other fits were classified as significant. (f) Fraction of neurons tuned to each variable across mice (*colored lines*) and sessions per mouse (*colored dots*). A large fraction of neurons (>50%) were significantly modulated by both position and progression-to-choice. Same as **Fig. 2d** but expanded to show other model variables. (g) Fraction of neurons tuned only to position in TONE trials or only to progression-to-choice. (h) The MI of cells tuned only to position (*black*), or progression-to-choice (*blue*) as shown in (g). The MI for each group was ∼ two-fold higher for the significant variable. (i) The distribution of the ratio of the MI for position versus progression-to-choice by only selecting cells that were significantly modulated by both variables. The distribution was centered around 0, with a larger skew towards spatial position. (j) Scatter of the kernel strength of choice licks during TONE trials, versus spontaneous licks for all cells significantly tuned to licks. The significant correlation confirms that cells were co-modulated by spontaneous and choice licks (Pearson’s correlation), thus irrespective of whether mice were performing the task and whether the water reward was present. (k, l) Same as (j), but for choice versus home, and spontaneous versus home licks. A significant decorrelation was observed, confirming that cells tuned to choice or spontaneous licks did not fire during the home port licks (Pearson’s correlation). Note the negative correlation between choice lick and home port lick-related firing even though both port types yielded the same amount of water reward, again indicating that the presence or absence of water reward *per se* was not a critical variable in determining neuronal firing patterns. (m) PSTH of all cells significantly tuned to licks using the PGAM. Unlike **Extended Data Fig. 9d**, which selectively examined tone cells, this analysis identified cells that were tuned to any lick during the session. Separating their firing into distinct lick types shows distinct groups firing to the NT, Home, and TONE choice licks. Note the peak firing still occurs before the first detected lick. (n) Average PSTH of all cell groups shown in (m). *Colors* indicate lick types as in (m). The shifted and broader peak of NT and Homeport licks reflects a different type of lick response compared to the tone choice licks and could emerge from place cells with a field at the ends of the track.

**Extended Data Figure 4.**
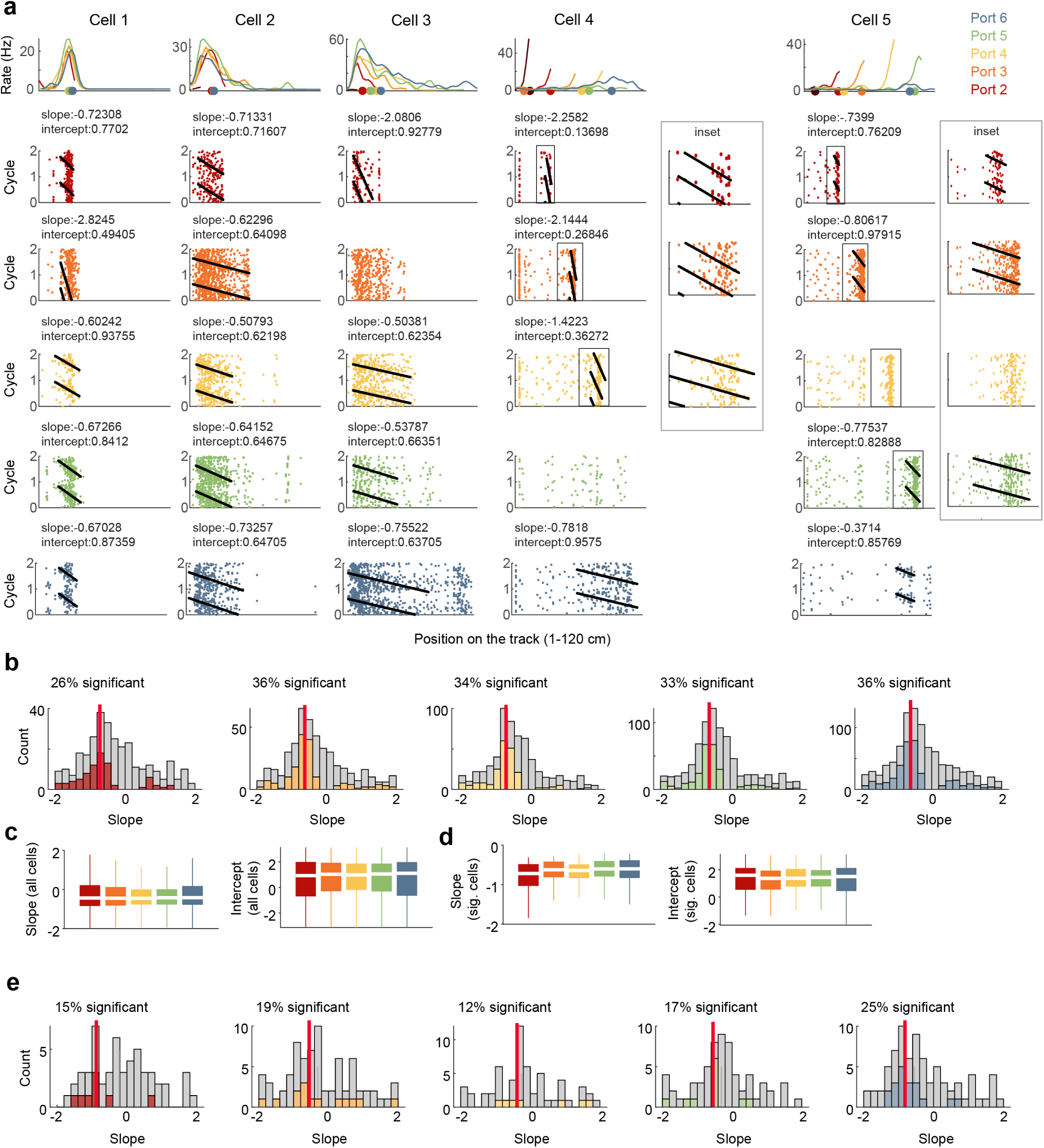
Phase precession of place and tone cells by approach to each port. (a) *Top row,* Five example cells across different sessions showing average ratemaps of the cell for each port. Cells 1-3, “place cells”, Cells 4-5, tone cells. *Colored dots on x-axis*, center of mass. *Rows 2-6*, theta phase by position plots for each cell, separated by trials to ports 2-6. For the tone cells, slopes were difficult to observe because of their small fields. Yet, clear examples of phase precession also occurred (see inset of ports 2-5 for Cells 4 and 5). The slope was calculated by identifying field boundaries for each port and scaling the place field size between 0 and 1. (b) *Left to right,* Distribution of phase precession slopes across ports 2-6 for place cells. *Gray bars,* all slopes. *Colored bars,* significant slopes. Colors as in (a). Fraction of significant slopes compared to all slopes is provided for each condition. *Red line,* median of significant slopes. (c) *Left,* Average phase precession slope of all place cells across port approaches. *Right*, Average phase precession intercept of all cells across port approaches. Colors as in (a). No effect was observed (n = 306-952 cells, Kruskal Wallis test followed by Tukey-Kramer post-hoc tests. Slope: Chi-sq (4,3097) = 1.42, *p = 0.84;* Intercept: Chi-sq (4,3097) = 0.55, *p = 0.97*). (d) *Left,* Average phase precession slopes of place cells with significant slopes across port approaches. *Right,* average intercept of significantly fit slopes (n = 80-344 cells, Kruskal Wallis test followed by Tukey-Kramer post-hoc tests. Slope: Chi-sq (4,1048) = 10.69, *p = 0.03;* Intercept: Chi-sq (4,1048) = 1.56, *p = 0.82*). Because the slope was calculated by normalizing place field size, no change in the slope implies that expanding fields would have spikes occurring in earlier theta phases at the same location compared to smaller fields (e.g., compare Port 2 and Port 6 trials of Cell 2 in (a). Despite similar slopes, the expanding field for Port 6 leads to a different theta phase of spikes). Colors as in (a). (e) Same as (b), but for tone cells. A smaller fraction of cells had significant slopes, possibly because of the smaller fields. Colors as in (a).

**Extended Data Figure 5.**
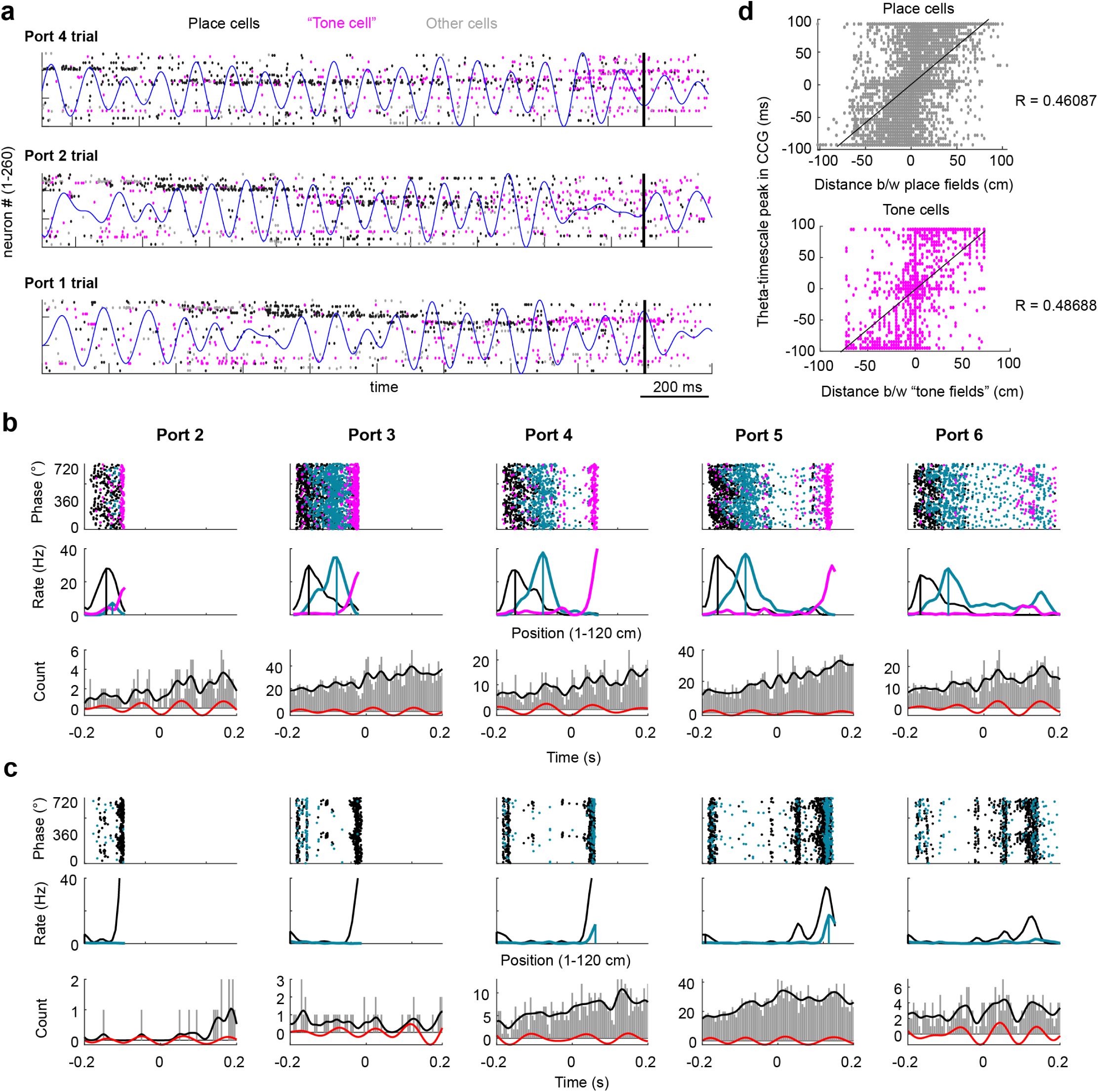
Theta sequences observed for both place and tone cells. (a) Spike rasters of three example trials from an example session (see **Fig. 4**). All active pyramidal cells are shown sorted using Rastermap (see experimental Procedures). *Black dots*, spikes from place cells, *pink dots,* spikes from tone cells, *gray dots*, spikes from non-classified cells. *Black vertical line* indicates time of first lick. *Blue trace*, filtered local field potential in the theta (6-12 Hz) band. Note that place cell sequences precede “tone cell” firing that ramps in the 0.5s preceding the lick. Note also that “tone cell” spikes are interleaved with place cell spikes and begin before the ramp preceding the lick. (b) *Top*, theta phase-of-spike by position plots for three example cells. *Blue and black*, place cells, *pink,* “tone cell”. *Middle*, Firing rate maps of each cell. Vertical lines, peak of the place fields. *Bottom,* Cross-correlogram (CCG) between the black and blue cells. *Red line*, filtered trace of the cross-correlogram in the theta-band. (c) Same as (b), but for two tone cells. (d) Distance to time compression. *Top*, scatter plot of the distance between place field peaks and the time lag in the CCG between cell pairs. *Bottom,* same as above but for tone cells. Distance between tone fields was calculated in the progression-to-choice domain. Both correlations were significant (Pearson correlation, Place cells: n = 5771 cell pairs, R = 0.4609, *1.8 x 10^-301^;* Tone cells: n = 2474 cell pairs, R = 0.4869, *p = 1.9 x 10^-147^*).

**Extended Data Figure 6.**
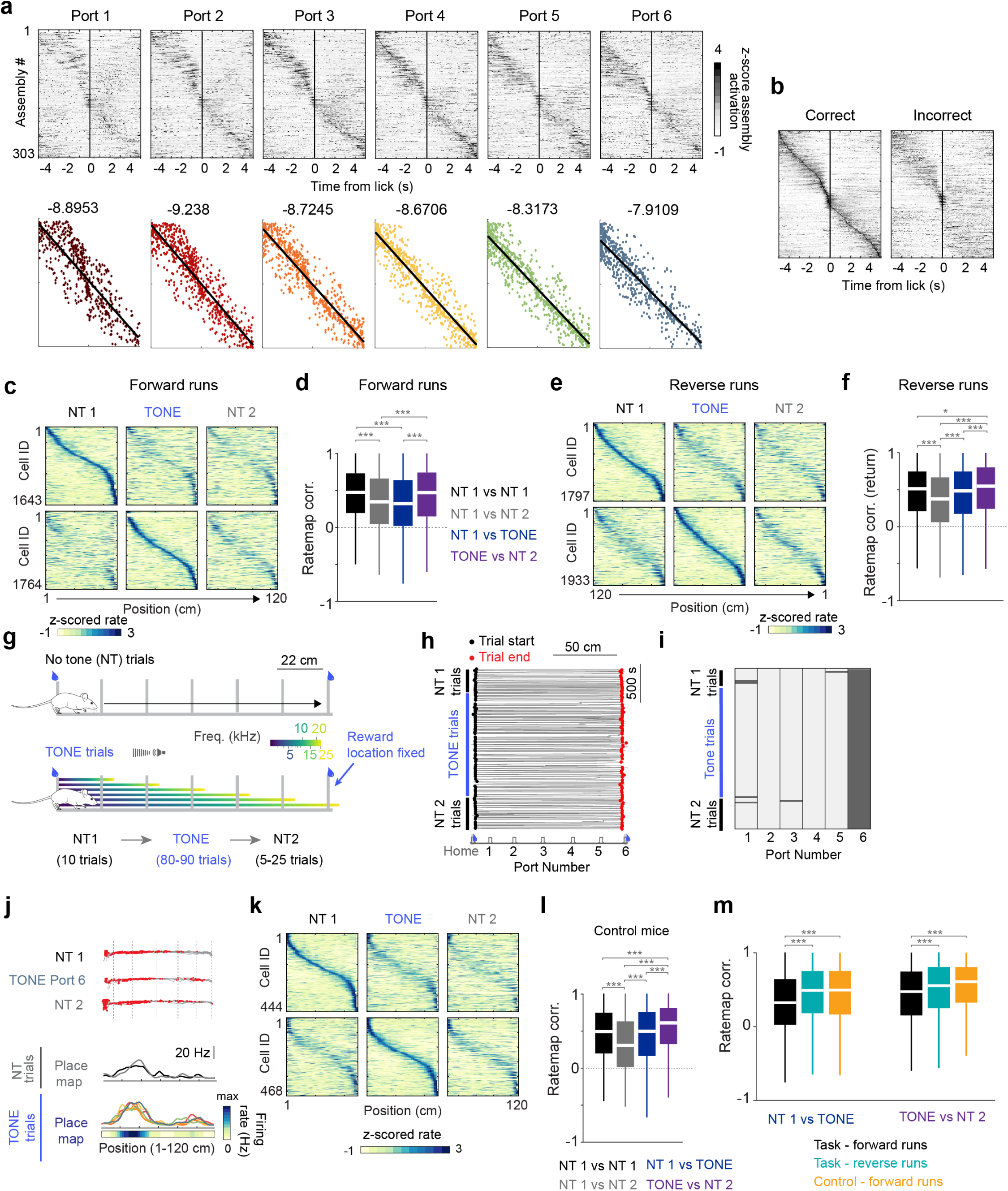
Evolution of cell assemblies and controls for spatial remapping. (a) Cell assemblies across all sessions were detected using ICA (see Experimental Procedures) and their activation around the end of the trial (i.e., lick at the chosen port) was calculated, separated by ports 1-6. *Top,* Assemblies were sorted by their activation profile calculated across all trials (see left plot in (b)) and the sorting order was maintained for the port-wise peri-event time histograms (PETH). The temporal width of the sequence expands from Ports 1-6. *Bottom,* Peak latency for each assembly as shown above. Sorting is maintained for Ports 1-6. *Lines and titles* show the regression slope decreasing from Ports 1-6. (b) Cell assembly PETH for the time of all correct choice licks. The same assembly sorting is maintained for error trial licks. Note similar cell assembly sequences in both correct and error trials in the 1s leading up to the lick. (c) Average firing rate maps for NT1, NT2 and TONE port 6 trials (to control for path length). *Top,* Cells with a detected place field (for place field definition, see Methods) in NT1 trials, sorted by their preferred location. *Bottom*, Same as above, but for cells with a place field during TONE trials. (d) Statistics for cell-by-cell correlations in (c) (n = 3004, 2767, 2869, 2618 cell-cell pairs; Kruskal Wallis test followed by Tukey-Kramer post-hoc tests, Chi-sq (3,11254) = 242.62, *p = 2.58 x 10^-52^*). (e) Same as (c) but for place fields on the return runs. (f) Same as (d) but for place cells on the return runs. There was a general drift in the spatial map, but no significant remapping between NT1 and TONE trial return runs (n = 2824, 2582, 2697, 2450 cell-cell pairs; Kruskal Wallis test followed by Tukey-Kramer post-hoc tests, Chi-sq (3,10549) = 199.99, *p = 4.22 x 10^-43^*; NT1 vs NT1 and NT1 and TONE Port 6, *p = 0.304*). (g) To control for the presence of the sensory cue, we trained a separate cohort of mice (n = 3) on a similar task. NT and closed-loop TONE trials alternated as in the main task, but rewards were always provided only at the two ends of the track. Thus, the tone had no significance. Tones would stop playing after 25 kHz. (h) Trajectory of a mouse from an example control session. (i) Heatmap showing licks detected at each port (*dark gray*) from the session in (h). (j) Example CA1 pyramidal cell recorded from a mouse trained on the control task. *Top,* Trajectories (*gray lines*) separated by NT1, TONE Port 6 and NT2. Spikes from a single neuron are overlaid in *red*. *Bottom,* Average spatial firing rates are shown for each trial type – NT1, individual TONE trials, NT2. *Rainbow shades*, TONE trial types 1-6. (k) Average rate maps from control mice for NT1, NT2 and port 6 TONE trials (to match the analysis performed for the auditory task mice, see (c)). *Top,* Cells with a detected place field in NT1 trials, sorted by their preferred location. *Bottom*, Same as above, but for cells with a place field during TONE trials. (l) Statistics for cell-by-cell correlations in (k). The results mirror the effect observed for the return runs in the auditory task mice, i.e., a general drift of the map led to a decorrelation between NT1 and NT2 trials (*gray*), but no significant remapping was observed between NT1 and TONE trials (*blue*) (n = 749, 674, 756, 676 cell-cell pairs; Kruskal Wallis test followed by Tukey-Kramer post-hoc tests, Chi-sq (3,2851) = 114.60, *p = 1.12 x 10^-24^*; NT1 vs NT1 and NT1 and TONE Port 6, *p = 0.974)*. (m) Comparison of the three conditions that are separated by similar time lags but show different magnitudes of remapping. Only the TONE forward runs led to remapping between NT and TONE trials, suggesting a significant role of task engagement that cannot be explained by changes in sensory inputs or representational drift (NT1 vs TONE, n = 2869, 2697, 756 cell-cell pairs; Kruskal Wallis test followed by Tukey-Kramer post-hoc tests, Chi-sq (2, 6319) = 174.49, *p = 1.28 x 10^-38^*; TONE vs NT2, n = 2618, 2450, 676 cell-cell pairs; Kruskal Wallis test followed by Tukey-Kramer post-hoc tests, Chi-sq (2, 5741) = 55.03, *p = 1.12 x 10^-12^*). *\*p<0.05, ***p<0.001.* All box plots show median ± interquartile; whiskers show range excluding outliers.

**Extended Data Figure 7.**
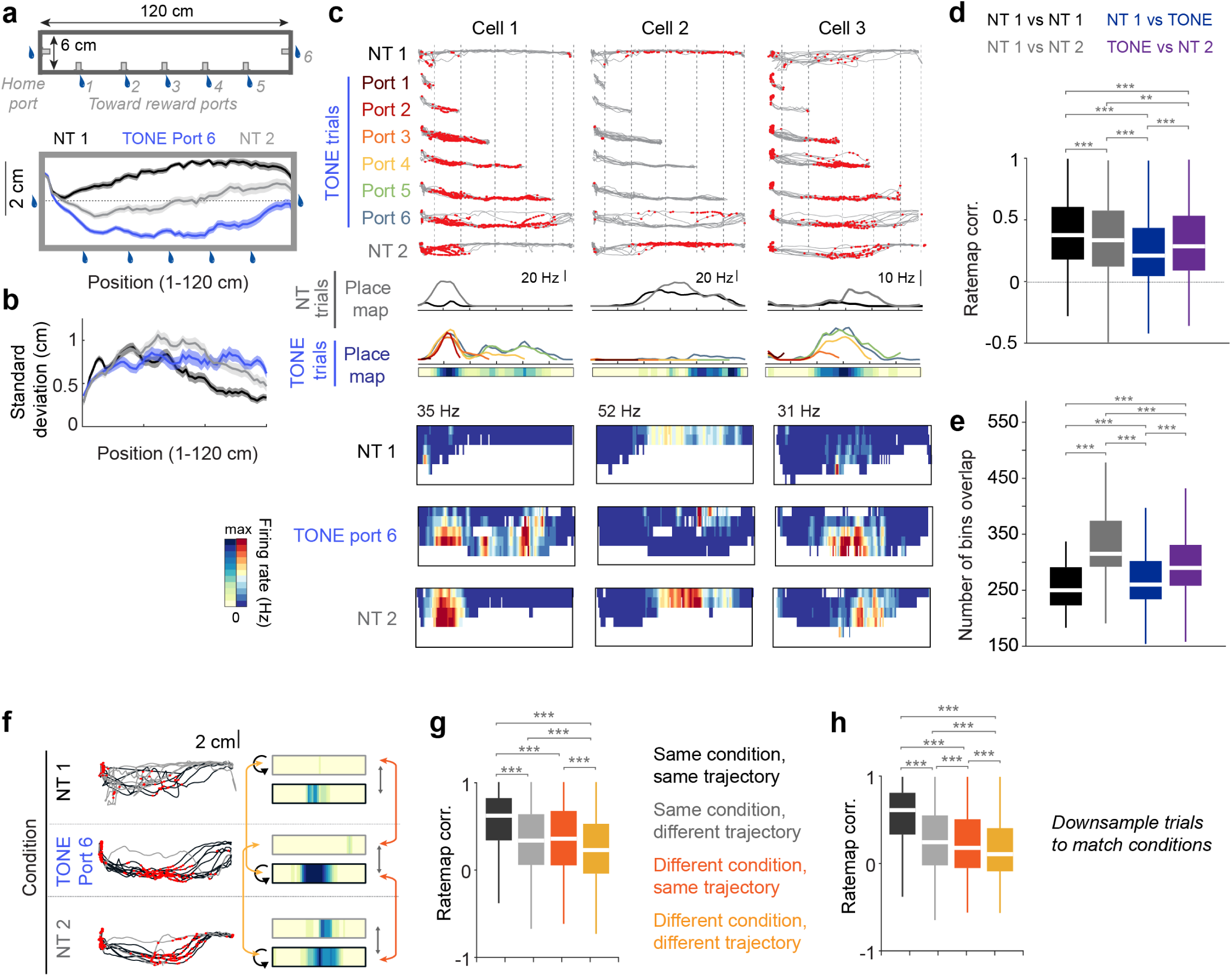
Spatial remapping depends on the mouse’s running trajectory. (a) *Top*, top-view schematic of the track showing the wall with the middle reward ports. *Bottom,* Movement trajectory (mean ± sem) across all mice for NT1 (*black*), TONE port 6 (*blue*) and NT2 trials (*gray*). The length and width ratios are distorted to emphasize small wall-guided deviation from a straight line on the track (dashed thin line). Note how mice avoided the port wall in NT1 trials, ran along the port wall in TONE trials, and midway between the two during NT2 trials. (b) Average standard deviation of the trajectory of the mouse along the width of the track. Trajectories were more variable for NT2, and TONE trials compared to NT1. *Colors* as in (a). (c) *Top*, three example cells with a place field recorded as mice ran with varying movement trajectories during NT1, TONE port 6 and NT2 trials. The firing of these cells was modulated by movement trajectory. *Middle*, one-dimensional linear maps for NT and TONE trials. *Bottom,* two-dimensional maps constructed for the same cells using 1 cm wide bins along the width of the track. Areas with no occupancy are shown in white. (d) Cell-by-cell correlation of the two-dimensional maps shown in (c). The decorrelation between NT1 and TONE trials persisted despite using a two-dimensional map (n = 1798, 1117, 2035, 2211 cell-cell pairs; Kruskal Wallis test followed by Tukey-Kramer post-hoc tests, Chi-sq (3,7157) = 256.57, *p = 2.48 x 10^-55^*). Only overlapping bins were considered for the correlation. (e) Average number of bins included for each correlation in (d). While the bin numbers were different, the trends did not match that of (d), suggesting the effect was not driven by the number of bins (Kruskal Wallis test followed by Tukey-Kramer post-hoc tests, Chi-sq (3,7163) = 1196.1, *p = 5.27 x 10^-259^*). (f) *Left,* spikes from an example cell (*red dots*) overlaid with the mouse’s trajectories during NT1, TONE port 6 and NT2 trials. Trajectories were clustered according to their distance from the top wall of the track (*gray versus black lines;* see Experimental Procedures). It was possible to have all trajectories within a condition to be of a certain cluster. *Right*, Rate maps were calculated independently for gray and black trajectories during NT1, TONE port 6, and NT2 trials (*box color* outlining the rate maps). Correlations between these rate maps were performed as depicted by the colored arrows. (g) Average cell by-cell rate map correlations separated by different conditions (NT1, TONE Port 6, NT2) and trajectory (*gray/ black*) comparisons as shown in (f). *Black,* within condition, the same trajectory was used as a baseline. *Gray,* changing trajectories even within the same condition altered the spatial map. *Orange*, the same trajectory, but different conditions also led to remapping. *Yellow*, there was an additive effect of trajectory and condition, suggesting both features combine to impact spatial firing (n = 5706, 4680, 10618, 10491 cell-cell pairs after pooling across similar groups; Kruskal Wallis test followed by Tukey-Kramer post-hoc tests, Chi-sq (3,31491) = 2425.1, *p = 0*). (h) Same as (g), but rate maps were down-sampled to match the number of black and gray trajectories. Only conditions which had at least one of each trajectory type were compared (n = 5706, 4601, 3366, 3362 cell-cell pairs after pooling across similar groups; Kruskal Wallis test followed by Tukey-Kramer post-hoc tests, Chi-sq (3,17034) = 2796.3, *p = 0*). Note that remapping occurred between NT and TONE trials even when the trajectories were the same. *\*\*p<0.01, ***p<0.001.* All box plots show median ± interquartile; whiskers show range excluding outliers.

**Extended Data Figure 8.**
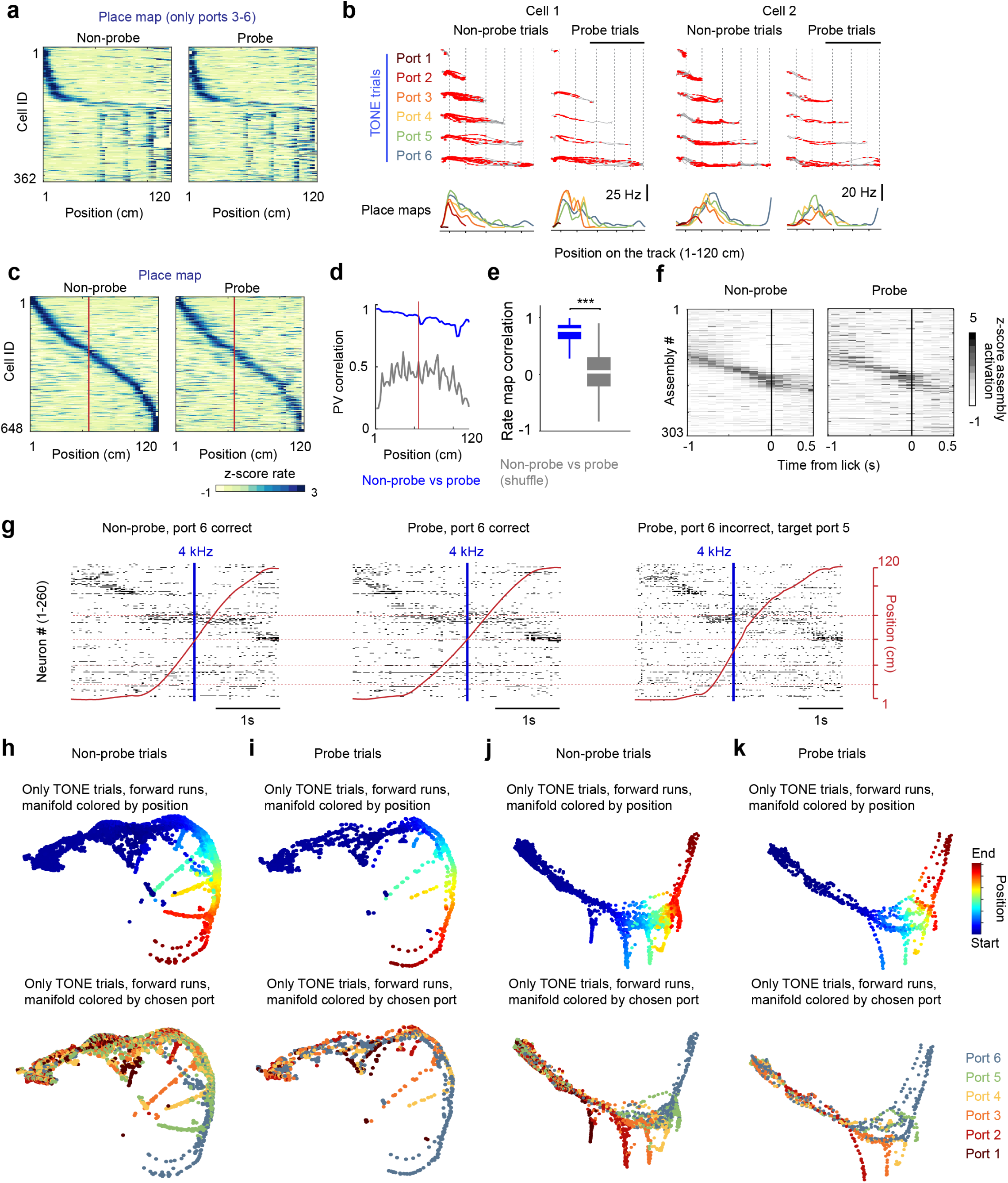
Similar evolution of cell assemblies during non-probe and probe trials. (a) Spatial maps of tone cells during non-probe and probe trials (sorted as **Fig. 3b**). Because no behavioral impairment was observed for ports 1-2 (**Extended Data Fig. 1g**), probe trials were only performed for ports 3-6 as targets. (b) Rate maps of two example place fields separated by port trajectory for probe and non-probe trials. A similar modulation of place fields by approach to the reward (**Extended Data Fig. 2h**) was also observed for probe trials. (c) Firing rate maps of all place cells detected using TONE Port 6 trials. The cells were sorted by their place field preference during non-probe trials and the same sorting was maintained for probe trials. *Red line* indicates position where tones were truncated during probe trials. Truncation of the tone did not affect the spatial map. (d) Population vector (PV) correlation for the heatmaps in panel c (*blue line*) compared to shuffled data (*gray*). The *red line* indicates the position where tones stopped during probe trials (as in c). (e) Cell-by-cell rate map correlations for the place cells in (c) (*blue*) compared to shuffled data (*gray*). Only cells with a place field after the red line, i.e., after the tones stopped playing during probe trials, were included in the analysis. Place maps remained stable, indicating that the removal of the tone did not impact the spatial map (n = 395 cells from 8 sessions in 2 mice, Wilcoxon rank sum test against shuffled data, z = 29.66, *p = 2.63 x 10^-193^*). (f) Cell assemblies across all sessions were detected using ICA (see Experimental Procedures) and their activation around the end of the trial (i.e., lick at the chosen port) was calculated. Cells are sorted by their latency in non-probe trials, and the sorting is maintained for probe trials. A stable cell assembly sequence persisted during probe trials, particularly in the ∼1 s preceding the lick. (g) Spikes sorted by rastermap (see Experimental Procedures) for the same session as **Fig. 4**. Three port 6 trials are shown: one non-probe (left), one probe correct (middle), and one probe incorrect (*right*). *Red line* is the mouse’s spatial position and blue line indicates the time when the tone was truncated (middle and right) or when the tone was playing at the probe frequency (*left*). *Red horizontal dashed lines*, port locations. Similar evolution of spiking activity was observed across all conditions. (h) Neural manifold generated using UMAP (see **Fig. 4** and **Extended Data Fig. 11-13**) showing non-probe TONE trials in an example session. Dots are individual time bins colored by position (*top*), or by trials according to the chosen lick port (*bottom*). (i) Same session as in (h), but for probe trials. The manifold shape is maintained, supporting the analysis that truncation of tones did not impact cell assembly sequences. Note no Port 5 probe trials occurred in this session. (j-k) Same as (h-i), but from a different animal. *\*\*\*p<0.001.* All box plots show median ± interquartile; whiskers show range excluding outliers.

**Extended Data Figure 9.**
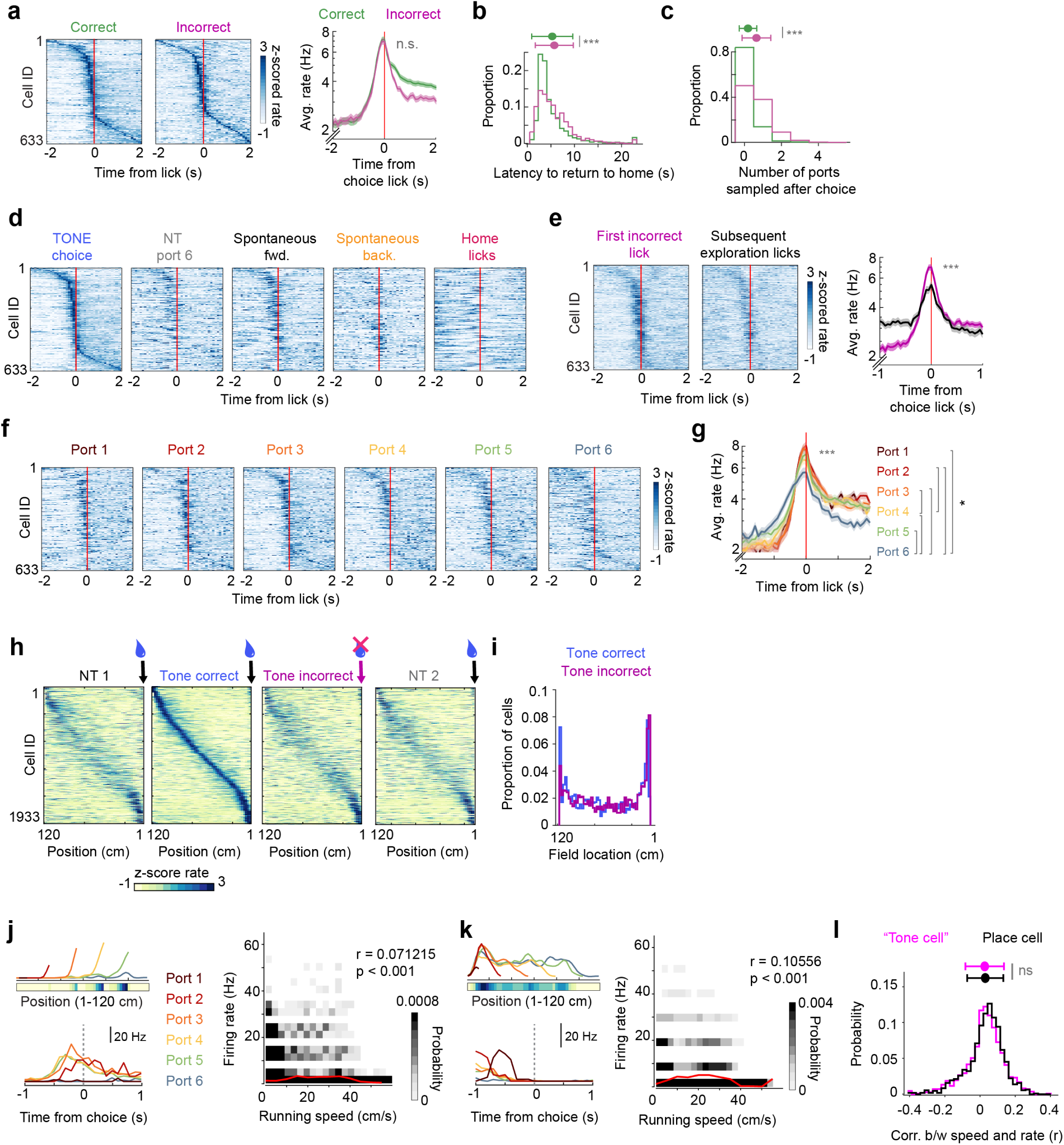
Tone cells were not directly modulated by reward or running speed. (a) *Left*, Peri-stimulus time histogram (PSTH) of tone cells for all choice licks during TONE trials, separated by correct *(rewarded)* and incorrect *(unrewarded error trial)* licks. To specifically examine the response to rewards, only tone cells firing around the target frequency 18-22 kHz have been included. Cells are sorted by their latency in the TONE correct trials and the sorting is maintained. 0s = first lick. *Right*, Average PSTH across tone cells for correct and incorrect licks. No significant difference was observed between the two (n = 633 reward-related tone cells, Wilcoxon signed rank test, z = 1.03, *p = 0.30*). (b) The latency to return to the home port after an incorrect lick was significantly higher than after a correct lick (n= 2079 vs 791 trials, Wilcoxon rank sum test, Z = -4.55, *p = 5.47 x 10^-6^*). (c) The number of ports sampled before returning to the home port after an incorrect lick was higher compared to a correct lick (Wilcoxon rank sum test, Z = -18.88, *p = 1.61 x 10^-79^*). (d) PSTH of all tone cells, separated by the distinct lick types as described in **Fig. 3 i, j**. Neurons are sorted by their latency in the TONE choice trials and the sorting is maintained. To specifically examine the response to rewards, only tone cells firing around the target frequency 18-22 kHz have been included. (e) *Left,* PSTH of tone cells around the chosen lick for all incorrect trials. *Middle,* PSTH of the same cells for subsequent exploratory samples after the incorrect lick. Sorting is performed as in (d). *Right*, the average firing rate of the cells for incorrect licks was higher than that for exploratory licks, suggesting some modulation by expectation of the outcome (n = 633 reward-related tone cells, Wilcoxon signed rank test, z = 6.8, *p = 1.03 x 10^-11^*). (f) PSTH of tone cells, separated by choice licks at Ports 1-6. Sorting is performed as in (d). Earlier ports showed a more robust response. Also see **Fig. 3m**. (g) Average PSTH of reward-related tone cells for choice licks during TONE trials, separated by port numbers (*rainbow colors*). Later (i.e., more probability) ports had lower firing rate modulation. Statistical comparison is between the average firing rate of the cells within ± 0.3 s around the lick (n = 633 cells from 6 mice, Friedman test followed by Tukey-Kramer post-hoc tests, Chi-sq (5,3160) = 77.96, *p = 2.24 x 10^-15^*). Post-hoc comparisons are shown on the *right*. Y axis is on a log scale. (h) Rate maps of all place cells during TONE trial port 6 return runs, shown for NT and TONE trials and separated by correct and incorrect trials. Cells are sorted by TONE correct trials. During incorrect trials, mice did not receive a water reward at the home port. Yet, the firing patterns remained the same. (i) Histogram showing the proportion of place fields at different positions on the track for the return runs. No difference was observed between incorrect and correct trials, even at the home port end of the track. (j, k) Two examples of CA1 tone cells. *Left,* Spatial map and PSTH of the cell for the first lick detected for each port. *Right,* Running speed versus binned firing rate for each cell, plotted as a probability distribution heatmap. *Red line*, mean firing rate by different speed. The correlation values for both cells were low (r, Pearson correlation). (l) Distribution of the correlation coefficient of speed and instantaneous firing rate for spatial (*black*) and tone cells (*pink*). The two populations were completely overlapping, indicating no preferential speed correlation with either cell “type” (n = 1129 tone cells and 1085 neurons with place fields from 6 mice, Wilcoxon rank sum test, z = -1.14, *p = 0.2531*. *ns, not significant*) *p<0.05, ***p<0.001. Line plots show mean ± sem.

**Extended Data Figure 10.**
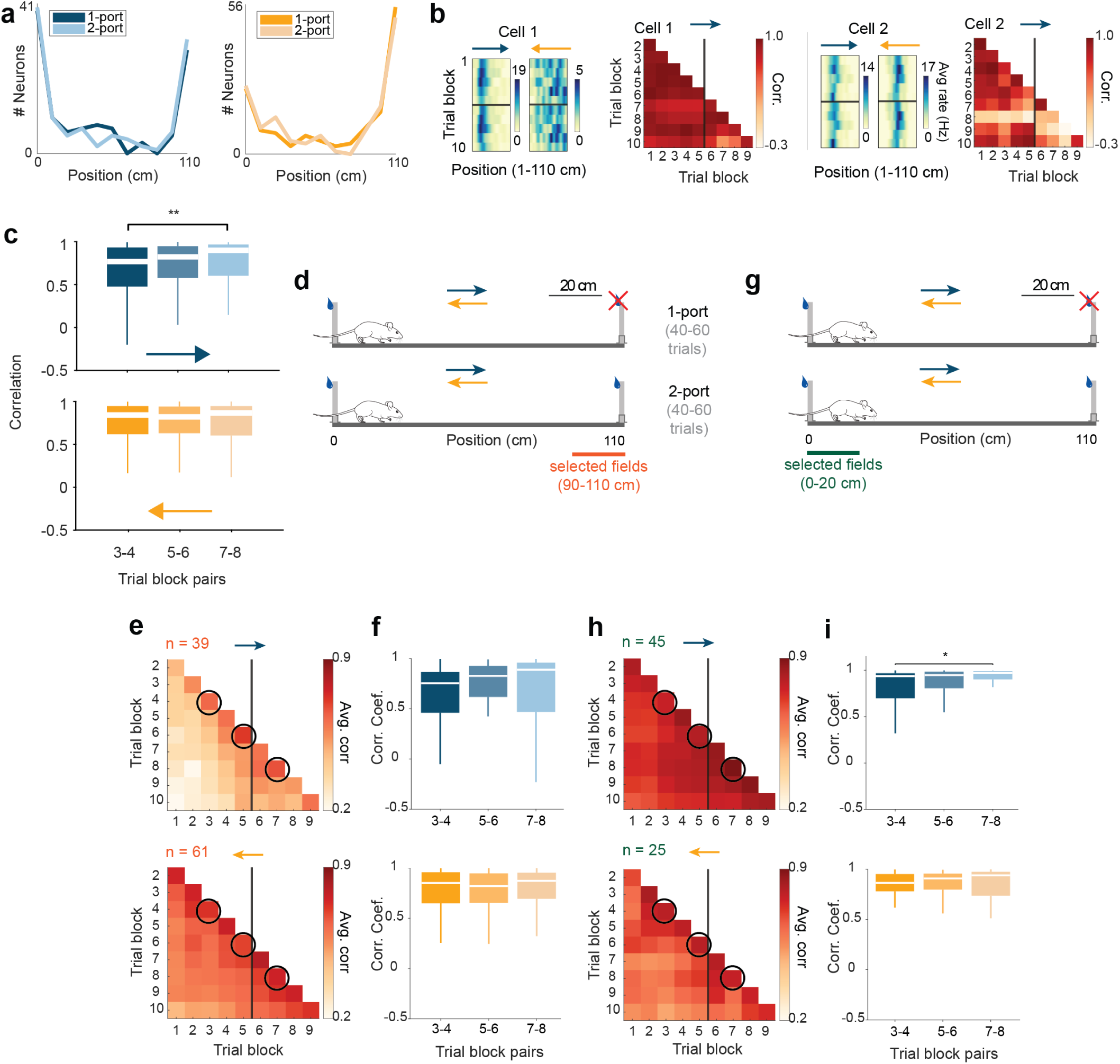
Firing rate maps do not change or overrepresent a new reward. (a) Distribution of place field peak locations (as in **Fig. 3e, f**) as the mice were running toward the changing reward port (*left*, n = 115 place cells) or when running away from the changing reward port (*right,* n = 146 place cells). Overrepresentation around the ports was observed for both the 1-port and 2-port conditions. (b) Trial block correlation for two example CA1 pyramidal cells. The *black line* indicates that the condition change happened between the 5^th^ and the 6^th^ trial blocks. (c) Average cell-by-cell correlation between blocks 3-4, blocks 5-6 and blocks 7-8 for all place cells (see **Fig. 3g**) highlights no significant change in the spatial map upon introduction of the reward (Friedman test followed by Tukey-Kramer post hoc tests, *Top* (to variable reward), n = 115 cells, Chi-sq (2, 222) = 9.16, *p* = *0.01; Bottom* (from variable reward), n = 146 cells, Chi-sq (2, 286) = 1.85, *p* = *0.397*). Significant post-hoc effect shows an increase in stability from blocks 3-4 and 7-8, and is not related to blocks 5-6, which is where the reward is introduced. (d) Schematic highlighting the selection criterion for neurons whose place field center was within 20 cm of the changing reward port. (e) Average cell-by-cell trial block correlation for neurons selected according to (d). *Black line* indicates that the condition change happened between the 5^th^ and the 6^th^ trial blocks. Black circles, selected trial block comparisons shown in (f). (f) Average cell-by-cell correlation between blocks 3-4, blocks 5-6 and blocks 7-8 for each running direction (Friedman test followed by Tukey-Kramer post-hoc tests, top; Chi-sq (2, 37) = 2.00, *p* = 0.3679, bottom; Chi-sq (2, 61) = 3.18, *p* = 0.2039). (g) Schematic highlighting the selection criterion for neurons whose place field center was within 20 cm of the stable reward port. (h) Same as (d), but for cells selected according to the criterion in (g). (i) Same as in (f), but for neurons selected according to (g) (Friedman test followed by Tukey-Kramer post-hoc tests, top; Chi-sq (2, 45) = 10.00, *p* = 0.0067, bottom; Chi-sq (2, 24) = 0.0833, *p* = 0.9592). Significant post-hoc effect shows an increase in stability from blocks 3-4 and 7-8, and is not related to blocks 5-6, which is where the reward is introduced. \**p* < 0.05, \*\**p* < 0.01. All box plots show median ± interquartile; whiskers show range excluding outliers.

**Extended Data Figure 11.**
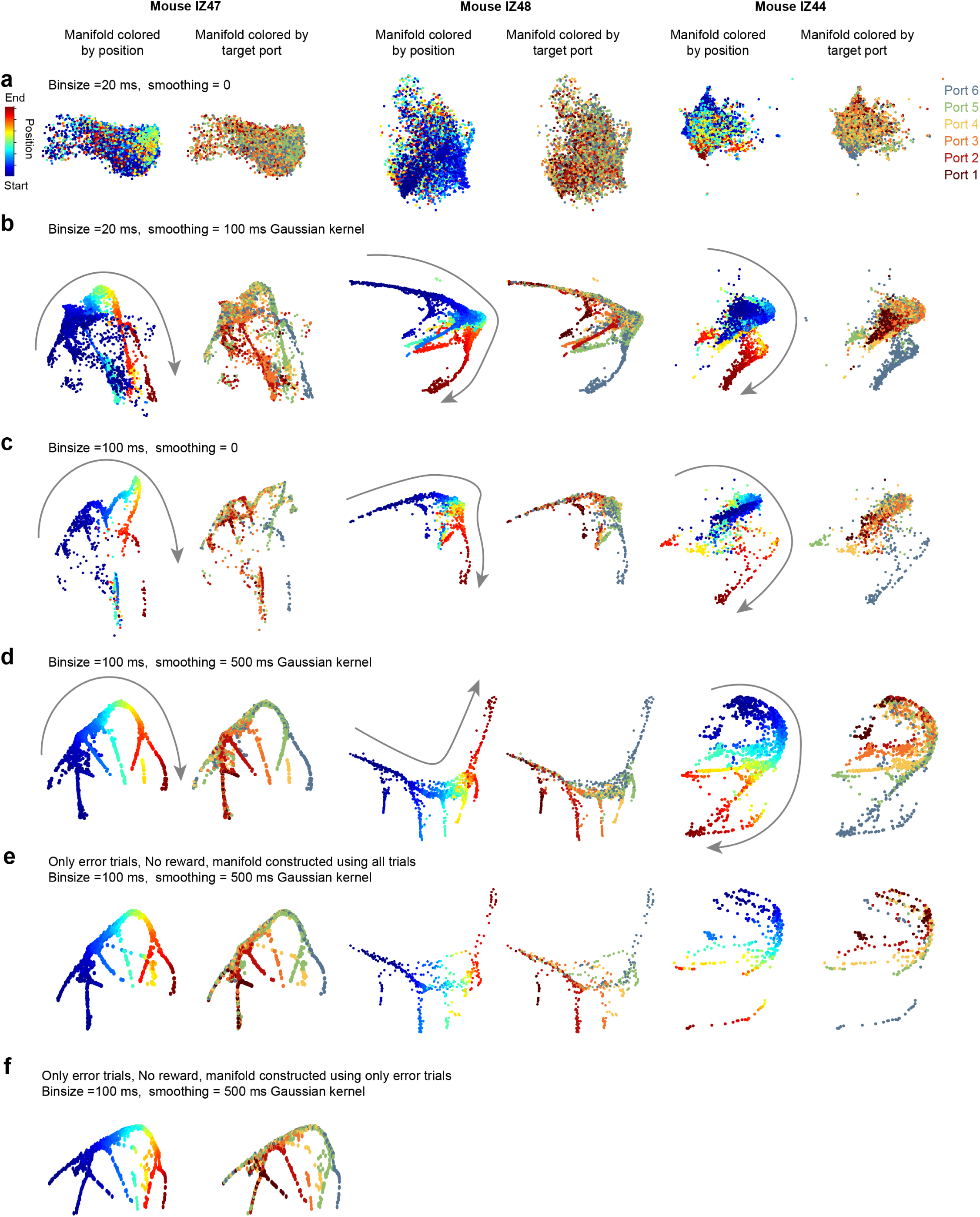
Effect of binning, smoothing and of rewards on the neural manifold. Neural manifold from three example sessions from separate mice – IZ47 (columns 1,2), IZ48 (columns 3,4) and IZ44 (columns 5,6). The activity of all neurons was binned into time bins of varying duration and various levels of smoothing and visualized on a lower dimension using UMAP. Each dot corresponds to a single time bin colored according to behavior variables. Only the forward running direction during correct TONE trials has been plotted. *Left*, Neural manifold colored by the spatial position of the mouse on the track. *Right*, Same manifold colored by the target port for that trial. Only times until the first detected lick on a trial are included. (a) Spikes were binned into 20 ms time bins and no smoothing was done. No consistent shape emerged for any of the sessions. (b) Spikes were binned into 20 ms time bins and smoothed using a 5 bin (100 ms) Gaussian kernel. (c) Spikes were binned into 100 ms time bins and no smoothing was performed. (d) Spikes were binned into 100 ms time bins and smoothed using a 5 bin (500 ms) Gaussian kernel. This condition resulted in the most consistent manifold shape across sessions, but branches were observed with lower binning and smoothing as well. (e) Same as (d) but showing only error trials. The manifold was constructed using the entire session. The manifold shape is conserved between correct and error trials, suggesting that the reward has no major role in the branching. (f) Neural manifold constructed from the session from IZ47, but only using error trials. No reward was ever delivered during these trials, yet the manifold looks almost identical to the one constructed using all trials.

**Extended Data Figure 12.**
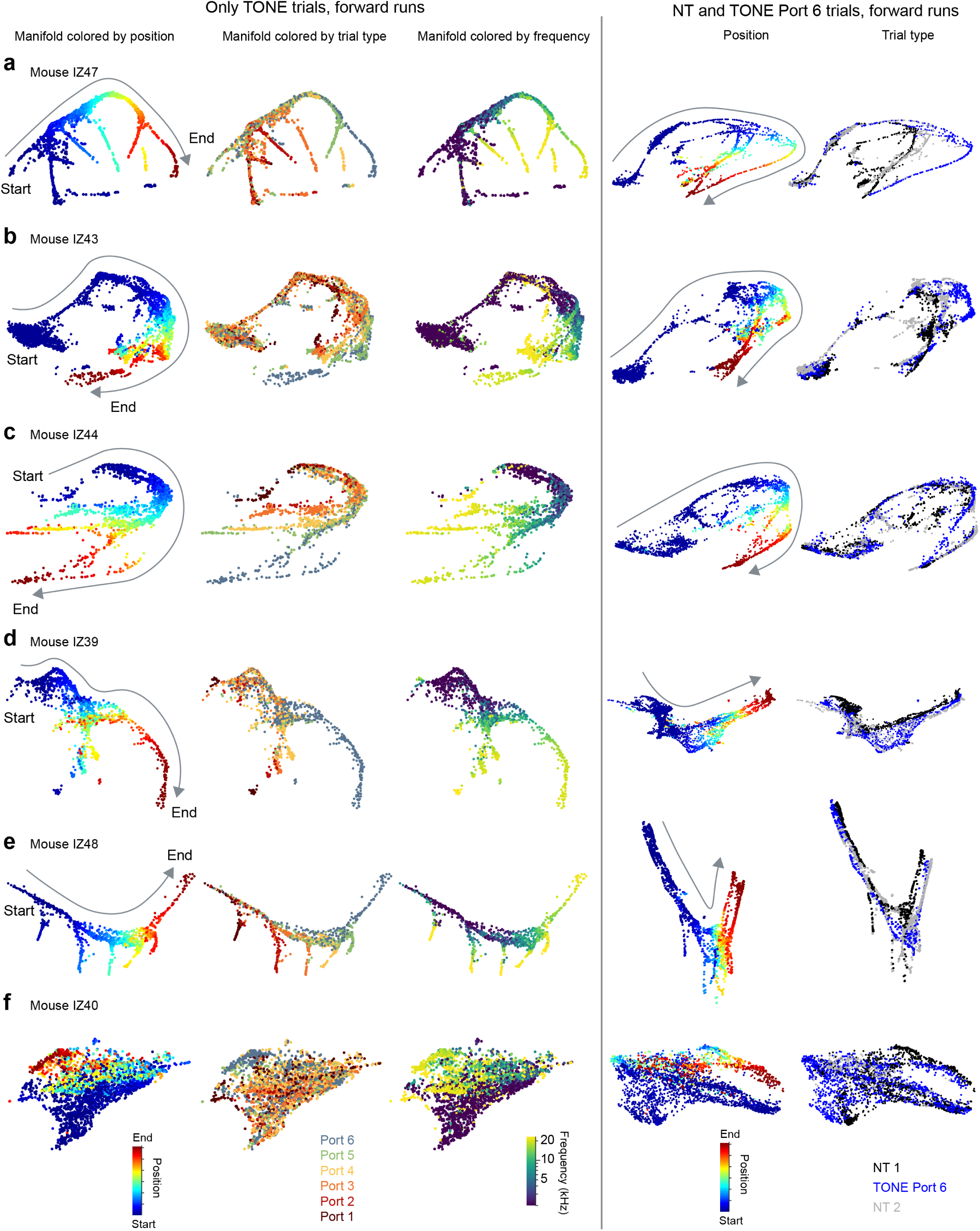
Distinct branches in the neural manifold were observed across several mice. (a-f) Neural manifolds from example sessions from six mice. Legend and color coding are shown in (f). *First, second and third columns.* Neural manifold that only shows the forward direction runs during correct TONE trials. *First column*, dots of the manifold are colored by spatial position. *Second column,* dots of the manifold are colored by the target port for that trial. Note that the manifold for each mouse had a similar shape with activity evolving along a common trajectory before diverging into distinct branches before the mouse chose a port. *Third column*, dots of the manifold are colored by the tone’s frequency. The divergence points occur roughly at around 10 kHz. *Fourth and fifth columns.* Neural manifold that only shows the forward direction runs during NT and TONE port 6 trials. *Fourth column*, dots of the manifold colored by spatial position. *Fifth column,* dots of the manifold colored by NT1, TONE Port 6, and NT2 trials. Neural trajectories differed depending on trial type – often resulting in more similar trajectories for Port 6 TONE and NT2 trials. Note that mouse IZ40 did not generate a neural manifold with any obvious structure, perhaps because of too few simultaneously recorded neurons (108 cells). See Extended Data Figure 13 for additional sessions and details.

**Extended Data Figure 13.**
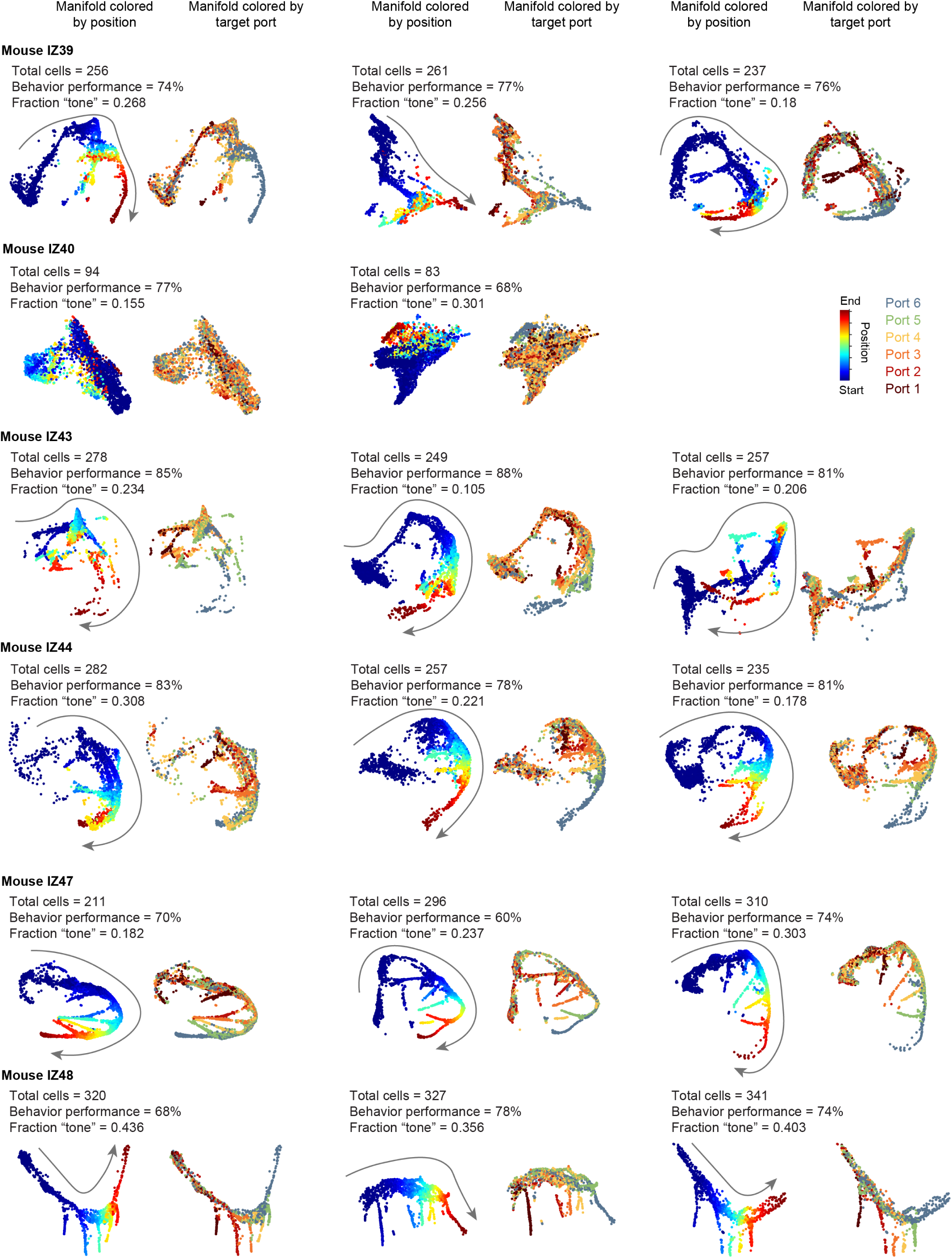
Variability in the neural manifold across sessions and animals. UMAP displays of 2 or 3 different sessions (*columns*) from each animal (*rows*). Only forward runs of correct TONE trials have been shown. *Left,* manifold colored by spatial position. *Right,* manifold colored by chosen port. The total number of recorded cells, behavior performance and fraction of tone cells are indicated for each session. While the number of cells was important to establish a general structure of the UMAP display (e.g., IZ 40), it did not appear to explain the precise shape of the UMAP. There was remarkable consistency across sessions within the same animal, yet similar features were also preserved across animals. Arrows show the direction from the start to the end of the track.

**Extended Data Figure 14.**
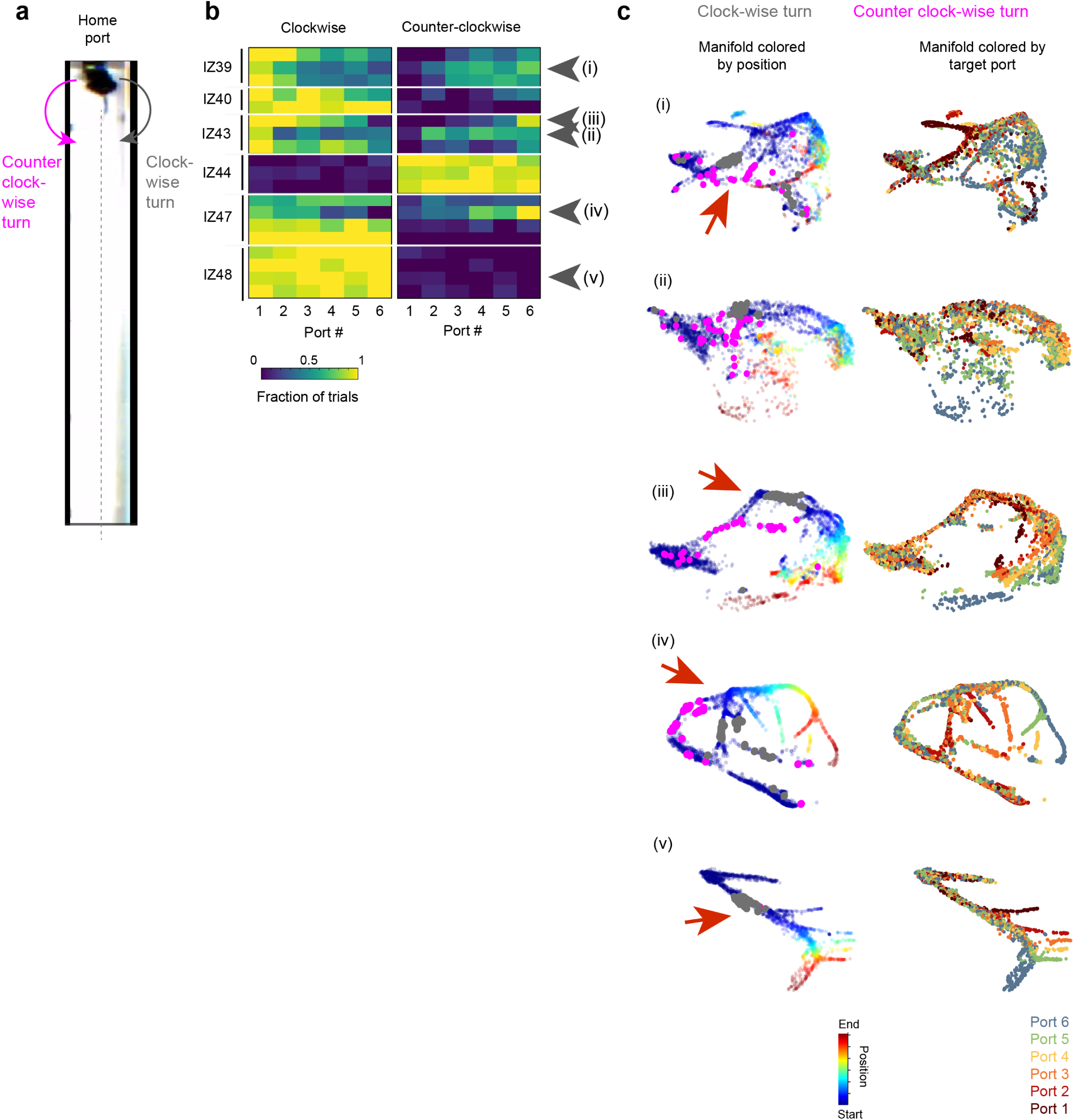
Direction of rotation of the animal is reflected in the neural manifold. (a) Top view schematic of the mouse licking at the home port. To start the next trial, the mouse can either turn clockwise (gray) or counter-clockwise (*pink*). (b) Fraction of clockwise versus counter-clockwise turns by session and animals. Some mice strongly preferred a single direction (e.g. gray arrowhead in v). In contrast, others showed a target port-dependent relationship in their preference to turn one direction or another (e.g., *gray arrowheads* i-iv). (c) Neural manifold colored by position and chosen lick port for the sessions marked by gray arrows i-v in (b). The manifold was rotated to optimize the view of the splitting branches (*red arrow*) for clock-wise and counter-clockwise turns (*Gray dots*, time of clock-wise turns, *Pink dots*, counter clock-wise turns).

**Extended Data Figure 15.**
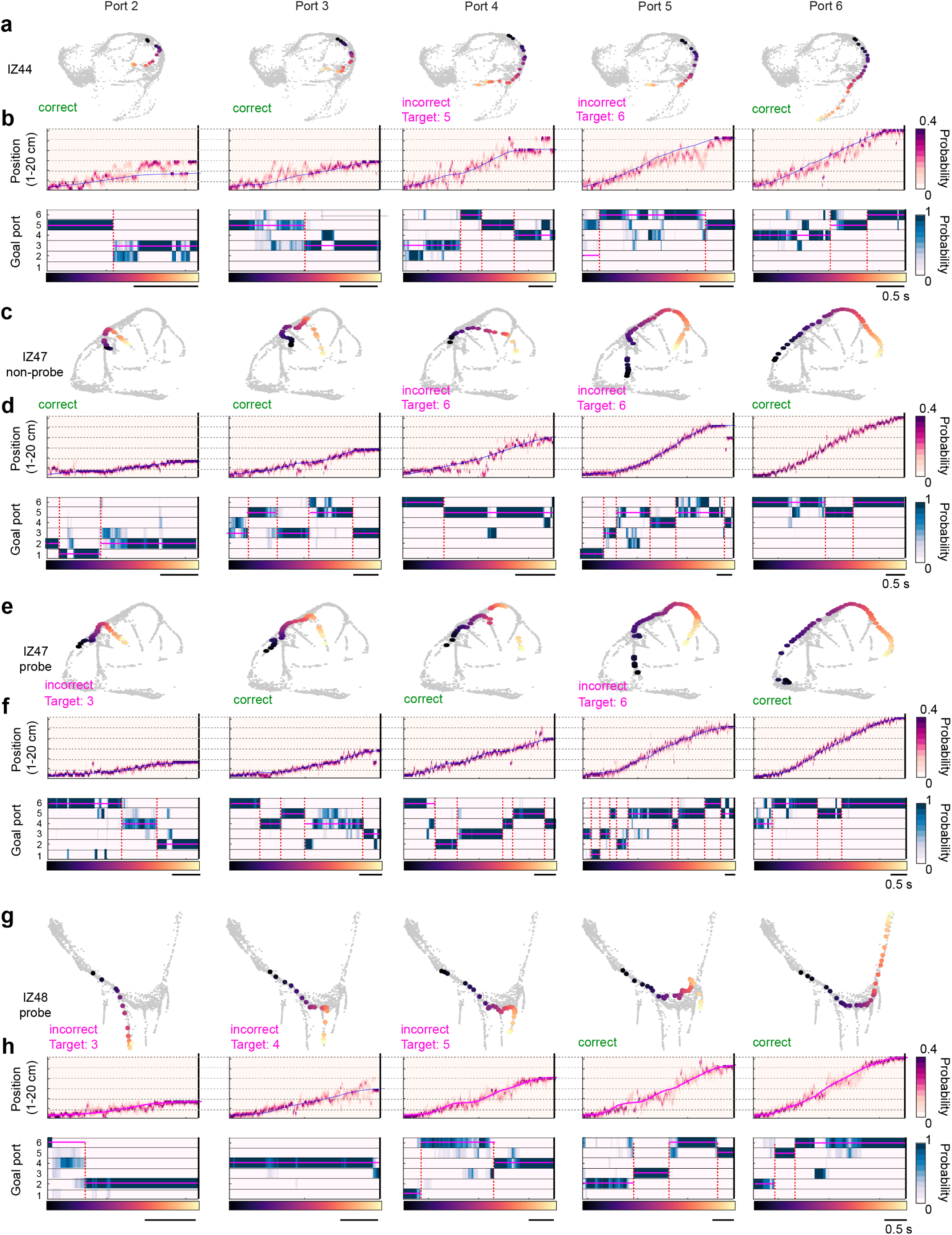
Examples of single trial UMAP displays, position and goal decoding across mice. (a) *Top*, UMAP (*gray dots*) of all forward TONE trials, overlaid with 5 different single trials going to target ports 2-6 (columns) (colors correspond to time in trial; colormap shown as x-axis in b). (b) 2-D spatial position and goal decoding (see Experimental Procedures) using 10 ms time bins. *Top*, Position decoding for the same trials as in (a). *Horizontal dashed lines*, port locations 1-6. *Blue line*, position of the animal. *Bottom*, goal decoding for the same trials s in (a). *Vertical black line* indicates the time of lick for that trial. *Vertical dashed red line* indicates significant change points (see Experimental Procedures) when the goal decoding switches. *Pink horizontal lines* show the mode goal decoded between two change points. (c, d) Same as (a,b) but for non-probe trials from a session by mouse IZ47. (e, f) Same as (c,d,) but for 4 kHz probe trials from the same session. The trajectory along the neural manifold, goal, and position decoding evolves similar to the non-probe trials. (g, h) Probe trials from a session from mouse IZ48 (same animal but different session as that of **Fig. 4**).

**Extended Data Figure 16.**
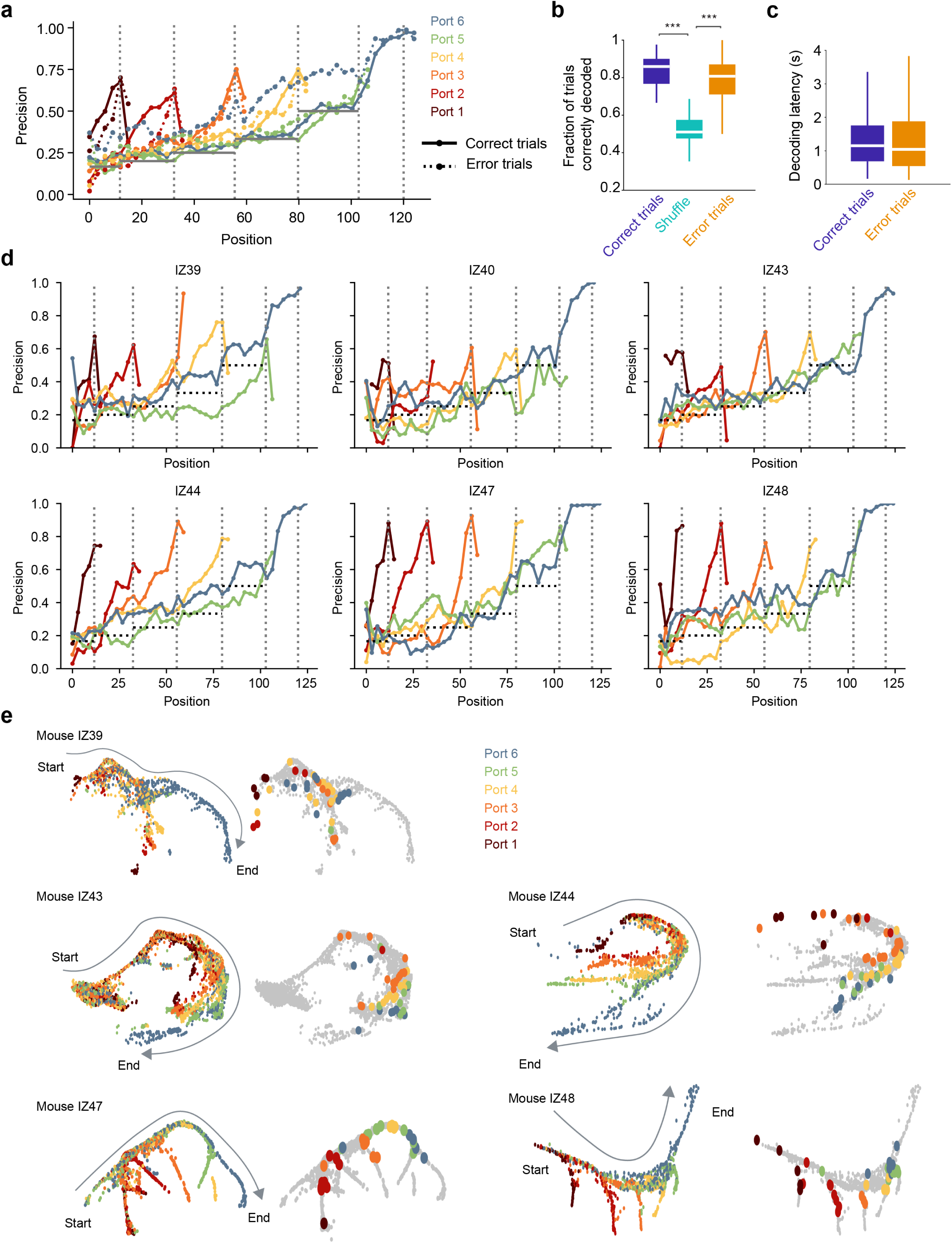
The target goal can be predicted from population activity. (a) Decoding of goal ports 1-6 (*colors red to blue*) from population spiking activity as a function of position. The decoder predicted the target port (*vertical gray dashed lines)* on both correct (solid lines) and error (dashed lines) trials above chance (*horizontal solid gray lines*) before the mouse reached the chosen port (37 sessions in 6 mice). (b) Using the change-point method (see Experimental Procedures), the final prediction of the decoded goal was estimated in every trial and compared to mouse’s choice. Note that both correct and error trials had high goal-decoding accuracy compared to a shuffle. In shuffle control the “choice” was randomly selected to be any of the remaining upcoming ports (n = 37 sessions from 6 mice, Friedman test followed by Tukey-Kramer post-hoc tests, Chi-sq (2,72) = 42.98, *p = 4.65 x 10^-10^*). (c) The time between the last change point in the decoder and the mouse’s choice of the upcoming port in correct and error trials, i.e., how much before the lick the target port could be predicted (∼1.1s; n = 1644, 519 trials, Wilcoxon rank sum test, z = 0.9, *p = 0.3676*). (d) Same as (a), but for individual animals (correct trials only). Mouse IZ40 with few cells had a poor decoding performance. *Horizontal dashed black lines* indicate chance levels. (e) Example neural manifold of each animal (except IZ40). All timepoints from forward TONE trials are shown. *Left*, dots colored by the target port. *Right*, all dots are shown in *gray*. Superimposed dots correspond to the time of the final detected change point during correct trials *Colors* indicate trials corresponding to ports 1-6. Decoding for each trial fell on the branch points for each target on the manifold, more clearly visible when the manifold branches were farther apart. Also see **Fig. 4f, g**.

**Extended Data Figure 17.**
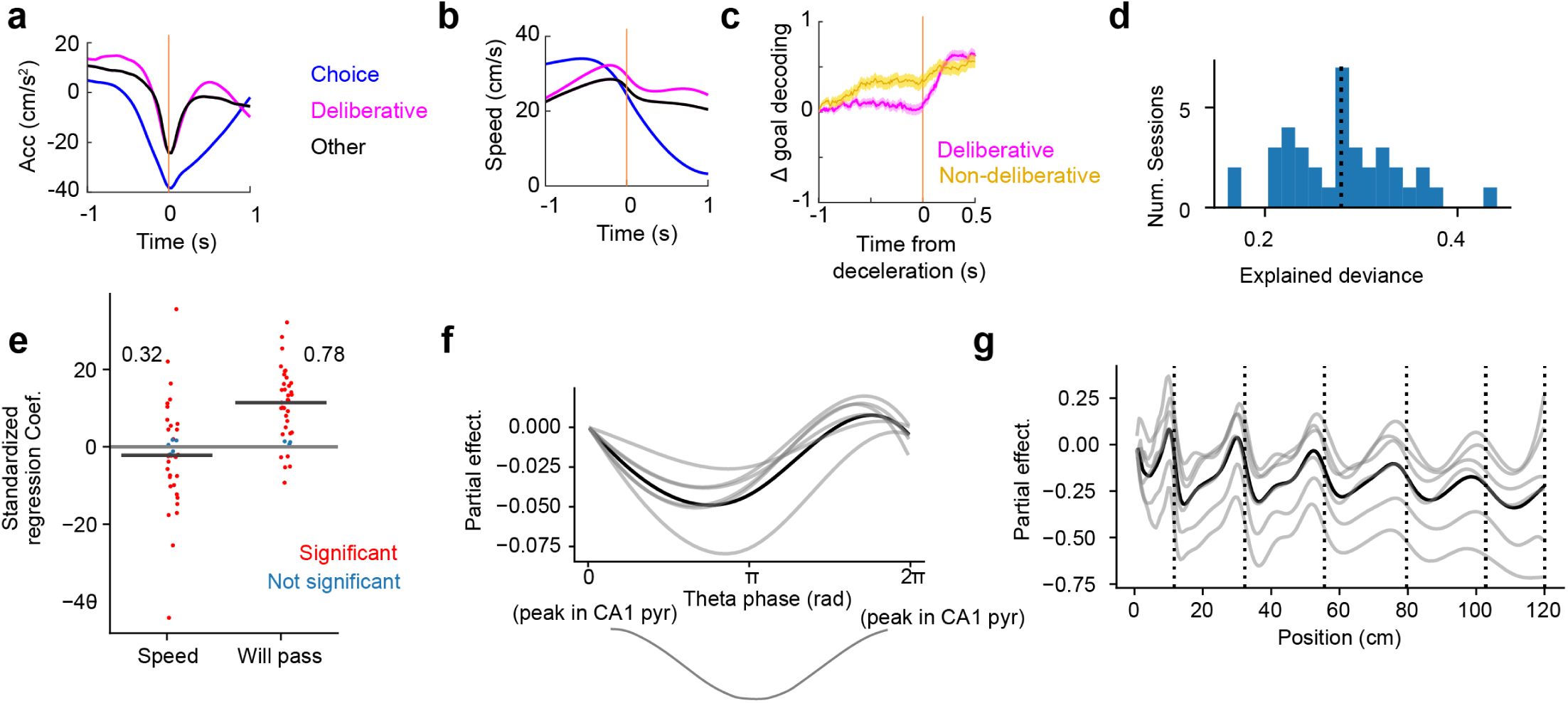
Relationship between speed deceleration and theta look-ahead within a trial. (a) All significant decelerations during forward TONE trials were separated into 3 categories – choice (approaching target port; *blue*), deliberative (slowing down near a non-target port, without licking; *pink*), and “other” (all other detected decelerations; *black*). Deliberative and other decelerations had similar acceleration profiles. Choice decelerations had a more pronounced trough. All sessions across all mice. Standard error is too small to be observable. (b) Same as (a) but showing the speed profile. Despite a significant deceleration, the speed of the mouse remained high during deliberative and other decelerations. (c) The change in goal decoding from the neuronal population against deliberative decelerations (*pink*, also shown in **Fig. 5g**). *Yellow*, as a control, we extracted random times when the mouse was crossing a port but did not decelerate (equal number of timepoints as deliberative decelerations). Thus, the jump in the decoder does not occur because of specific port cues. Lines show mean±SEM. (d) A generalized additive model (GAM; see Experimental Procedures) was fit to predict the summed posterior beyond the upcoming port, using the linear regressors *speed* and *will_pass* (whether the animal will pass the upcoming port) and smooth function (via B-spline basis functions) regressors theta phase and position. *Blue bars* are the distribution of explained deviance of the model in all sessions. The vertical dotted line marks the mean. (e) Standardized regression coefficients (per session) for the linear terms: *Speed* and *Will_pass,* to predict whether the position decoding posterior is beyond the upcoming port. Color marks whether the coefficient is significant, based on the asymptotic standard errors based on the Fisher information matrix. Numbers refer to the fraction of sessions with positive and significant regression coefficients. This plot shows that the running speed did not impact the position decoding posterior, which instead was strongly dependent on whether the mouse was going to run past the port or not. (f) The partial effect of theta-phase of the summed posterior after the upcoming port. Each *grey line* is the average over sessions within an animal. *Black line*, group average over all sessions. The *grey line* on the bottom is the LFP theta cycle. We observe an increase in the partial effect in the ascending, but not the descending phase of theta. Thus, there is a greater “lookahead” (or probability of the position decoding posterior to be beyond the upcoming port) specifically during the ascending phase of theta. (g) Similar to (f), but with the partial effect of *position*. The vertical dotted lines mark the positions of the ports. The partial effect ramps up towards the port and falls after the port by design, because the predicted variable is the summed posterior beyond the next port. It does not show there is more theta lookahead (relative to the animal) as it approaches the port. The term is only included as a control and the result confirms that the GAM fit is reasonable.

